# Chemiluminescent probes allow for the rapid identification of colibactin-producing bacteria

**DOI:** 10.64898/2026.01.02.697419

**Authors:** Miguel A. Aguilar Ramos, Sara Gutkin, Maya David, Doron Shabat, Emily P. Balskus

## Abstract

The *pks* (or *clb*) gene cluster, which produces the genotoxic natural product colibactin, is encoded by human gut *Enterobacteriaceae*, including many commensal strains of *E. coli*. Colibactin crosslinks DNA and is implicated in colorectal cancer development, highlighting the importance of identifying colibactin-producing gut bacteria within biological samples. In this study, we develop phenoxy-dioxetane chemiluminescent probes that selectively react with a critical colibactin biosynthetic enzyme, the serine peptidase ClbP. We show that these chemiluminescent probes have superior sensitivity, speed, and detection capabilities compared to previously reported fluorescent ClbP probes. Furthermore, we employ these chemiluminescent probes to detect *pks*^+^ *E. coli* directly in complex stool suspensions. These probes will enable multiple applications requiring detection of colibactin-producing bacteria, including the identification of ClbP inhibitors and the screening of clinical samples.

## Introduction

The human gut microbiome harbors hundreds of microbial species that have been implicated in both health and disease states, including colorectal cancer (CRC) development.^1^ Gut bacteria are associated with CRC in clinical studies and play causal roles in CRC tumorigenesis in animal models.^2–4^ One of the most prominent gut microbial factors linked to CRC is the *pks* (or *clb*) gene cluster, a biosynthetic pathway that produces the chemically unstable, genotoxic nonribosomal peptide-polyketide natural product colibactin.^5,6^ Colibactin is predicted to have a pseudodimeric structure containing two cyclopropane warheads that react with adenines, generating interstrand crosslinks in DNA,^7,8^ double-strand breaks, and genomic instability.^5,9–13^ Further, mutational signatures arising from colibactin exposure (SBS88, ID18) have been characterized in *in vitro* eukaryotic models^14,15^ and detected in CRC and other cancer genomes, indicating that humans are exposed to colibactin.^16^ These mutations have been identified in known CRC driver genes, including *APC* (adenomatous polyposis coli), a tumor suppressor gene that is commonly mutated in CRC. *APC* contains ID18, the characteristic indel mutational signature produced by colibactin, in 25% of its driver mutations, which suggests direct involvement of colibactin in cancer development.^14,15,17–19^

The strong associations between *pks*^*+*^ gut bacteria and CRC underscores a need to reliably detect colibactin production in complex clinical samples. However, chemical or analytical tools that can accurately and swiftly detect the presence of *pks*^*+*^ bacteria in complex matrices are not available. Methods to detect *pks*^*+*^ organisms that rely on amplification of genetic material, such as PCR or loop-mediated isothermal amplification (LAMP), are cumbersome in that they require the extraction and purification of nucleic acids, and do not confirm that biosynthetic genes are being expressed.^20^ Due to its chemical instability, colibactin has eluded conventional isolation and structural characterization, and is challenging to detect via standard methods such as LC–MS analysis.^21^ However, the reactivity of colibactin biosynthetic enzymes can be exploited for detection. Specifically, the final step of colibactin biosynthesis employs a self-protection mechanism involving the hydrolysis of two *N*-myristoyl-D-asparagine units (prodrug scaffolds) from an inactive biosynthetic precursor (precolibactin) by a periplasmic serine peptidase ClbP (**Figure 1A**).^22–24^ ClbP (and the rest of the *clb* genes) are expressed in a constitutive manner in *E. coli*, with expression occurring at very low levels in early growth stages and progressively increasing in the mid-log phase. *pks* gene cluster expression is also increased under certain conditions (iron deficiency, low oxygen, nutrient availability, sublethal amounts of antibiotics, etc.)^25^ ClbP selectively processes substrates containing the prodrug scaffold using an extensive hydrogen-bonding network to recognize the D-asparagine side chain and non-polar contacts with the hydrophobic myristoyl tail.^24^ The C-terminal end of the prodrug scaffold is amenable to modification, enabling the development of probes to detect ClbP activity. Fluorescence based–ClbP probes have been previously developed,^26,27^ consisting of coumarin fluorophores linked to the key *N*-acyl-D-asparagine ClbP recognition motif (**Figure 1B**). Upon hydrolysis by ClbP, the prodrug motif is released along with either a self-cleaving or an active coumarin, yielding detectable fluorescence. Probe optimization led to the exchange of the myristoyl chain for a 4-phenylbutyryl chain and the removal of the alanyl linker to achieve greater solubility and signal intensity.^26^ These fluorescent probes have been used to monitor ClbP activity in vitro and in bacterial culture and have enabled high-throughput screens for ClbP inhibitors.^28,29^ However, they have several disadvantages, including high autofluorescence in samples, low sensitivity and narrow dynamic range that limit their use in complex sample types. These limitations can be addressed through the design of chemiluminescent probes. In contrast to their fluorescent counterparts, luminescent probes do not require advanced op-tics to be detected.^30^ Further, chemiluminescent probes are more sensitive, as illumination and subsequent fluorophore excitement is not needed.^31^ The signal-to-noise ratio produced by chemiluminescent probes is significantly higher than their fluorescent counterparts,^32^ which is beneficial when working with complex biological matrices that present autofluorescence. Such chemiluminescent probes employ a modified Schaap’s luminophore, an adamantylidene-dioxetane based probe, which results in highly emissive luminescence in aqueous solutions when the dioxetane decays in aqueous solutions.^32^ Recent advances in chemiluminescent probe development have yielded probes that can selectively detect enzymes from the bacterial pathogens *Salmonella sp*., *Listeria monocytogenes, Staphylococcus aureus, Pseudomonas aeruginosa*, and *Mycobacterium tuberculosis*.^30,33,34^ Arrays of chemiluminescent probes for enzymatic activity have also been used to classify bacterial species using nearest neighbor algorithms.^35^ However, this strategy has not yet been applied to probes that detect colibactin production.

**Figure 1.**
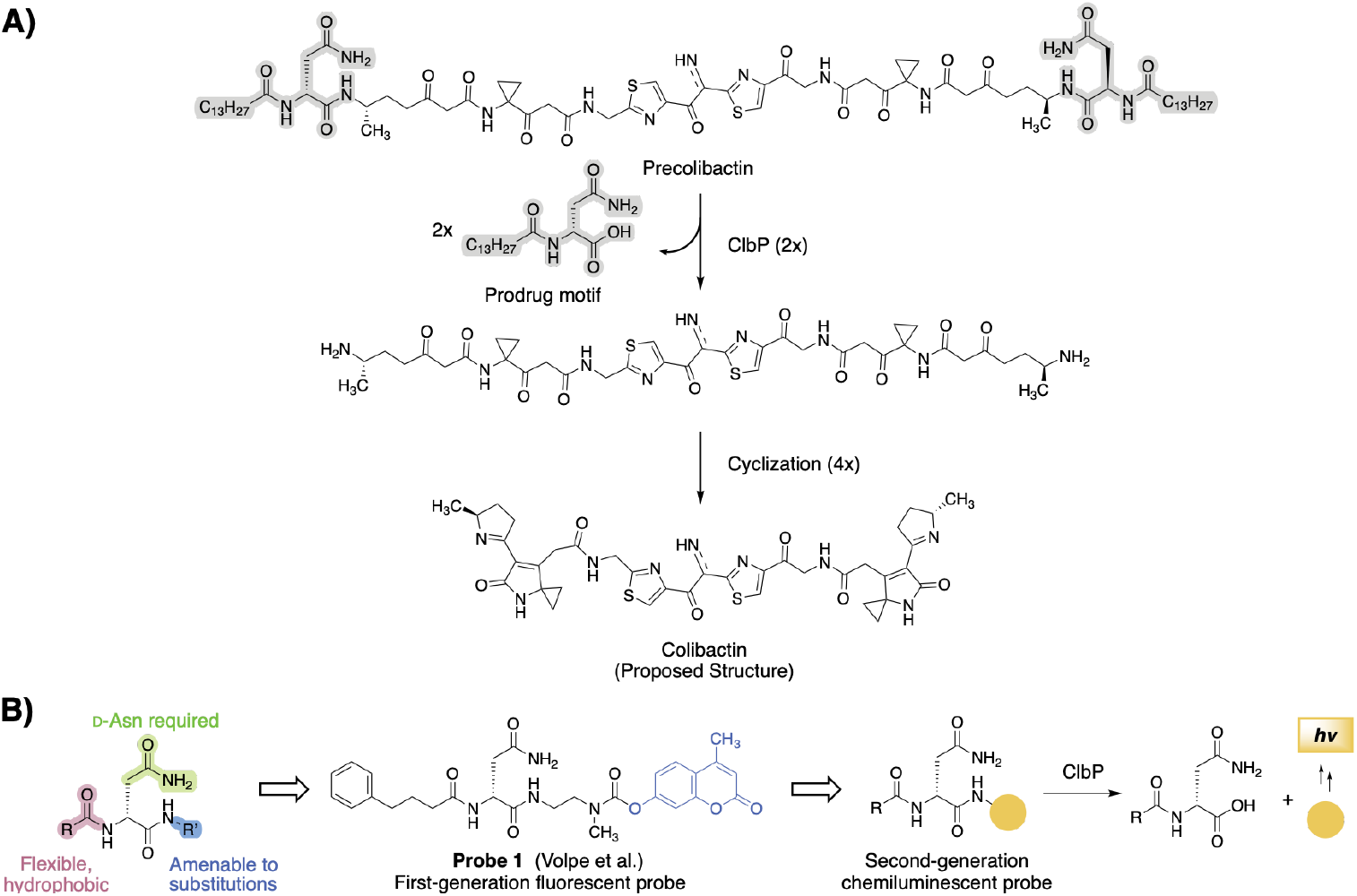
Design of a chemiluminescent probe for detection of the colibactin biosynthetic enzyme ClbP. A) The inactive biosynthetic precursor precolibactin is processed by ClbP to release the gut bacterial genotoxin colibactin. ClbP selectively recognizes substrates containing an N-acyl-D-Asn prodrug scaffold (highlighted in gray) B) Substrate preferences of ClbP guided the design of fluorescent activity-based probes and can be applied for second-generation chemiluminescent probes.

Here, we describe a set of chemiluminescent probes for ClbP activity that can rapidly and selectively detect *pks*^+^ gut bacteria. These probes can quantitatively detect signal from as few as 10^4^–10^5^ *E. coli* cells per mL in a one-hour measurement. Furthermore, the probes can detect *pks*^*+*^ gut bacteria in stool sample resuspensions, highlighting their utility and robustness in complex microbial communities. To our knowledge, this is the first example of an activity-based chemiluminescent probe suitable for detection of a specific bacterial enzyme in stool samples. Altogether, our results illustrate that chemiluminescent probes are promising candidates for detection of *pks*^+^ gut bacteria, with potential applications in CRC diagnosis and/or prevention. Furthermore, this study highlights the broader potential of chemiluminescent activity-based probes for detection of specific activities in the human gut microbiome.

## Results and Discussion

### Probe design and synthesis

We envisioned achieving selective and efficient processing of a chemiluminescent probe by ClbP using a design principle resembling that used in earlier fluorescent probe development (**Figure 1B, Figure S1**).^26,27^ Specifically, we devised a probe bearing the *N*-acyl-D-asparagine prodrug motif tethered to a self-immolative linker and a highly-emissive dioxetane (**Figure 2A**). An initial set of probes were designed to contain two different acyl chains (myristoyl and 4-phenylbutyryl) combined with either a free acid or a methyl ester, which is more likely to permeate the cell membrane, at the acrylate substituted luminophore (**Figure 2B**). Hydrolysis of these probes by ClbP would release the prodrug motif as well as trigger self-immolation of the linker. The dioxetane then undergoes chemiexcitation whereby the excited benzoate productively decays and emits green photons^32^. We synthesized these probes using the scheme outlined in **Figure 2C**. Briefly, we coupled Fmoc-d-Asn-OH with 4-aminobenzyl alcohol to form amide **5**, which was then treated with sodium iodide and trimethylsilyl chloride to form iodide **6. 6** was then coupled with the known dioxetane-bearing fragment **7**^36^ to afford enol ether **8**, a common precursor to all target probes. Acylation of **8** with the appropriate activated acyl succinimide and optional hydrolysis using lithium hydroxide afforded precursor enol ethers **9-12**. Treatment of these intermediates with singlet oxygen as previously reported afforded our desired probes **2-4**.

**Figure 2.**
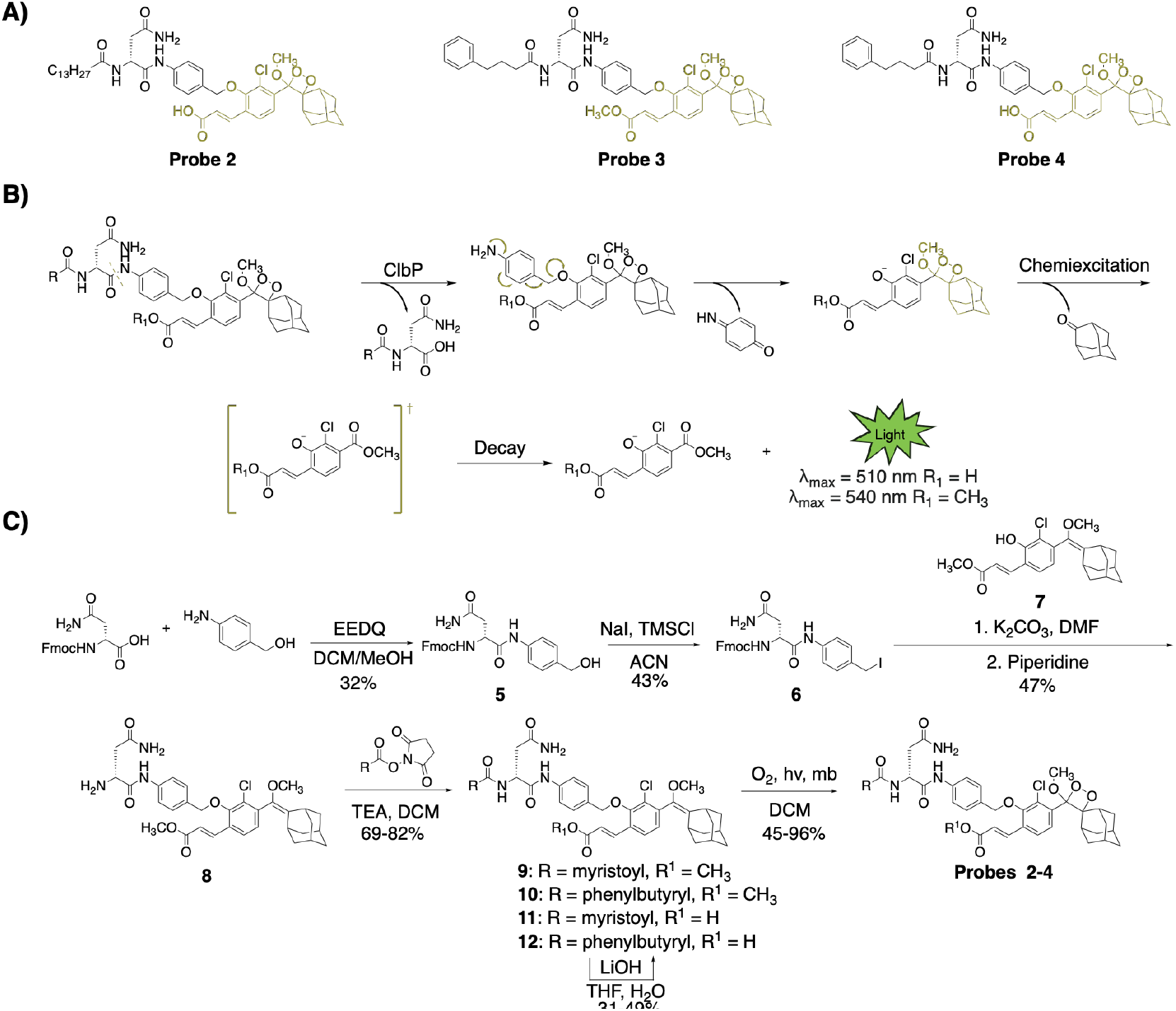
Design and synthesis of chemiluminescent probes for ClbP detection. A) Structures of chemiluminescent probes 2-4. B) Proposed mechanism of probe function. Following proteolytic cleavage, the probes self-eliminate, become chemiexcited, and release green light in aqueous solutions. C) Synthesis of chemiluminescent probes that target ClbP.

### In vitro testing of chemiluminescent ClbP probes

We next characterized the activity of the probes towards purified ClbP.^26^ Each probe was incubated individually with either wild-type (WT) 6xHis-C-tagged ClbP or an inactive ClbP mutant in which the active site serine is replaced by an alanine (ClbP S95A)^26^ (**Figure 3A**). All the tested probes yielded luminescence signals when incubated with WT ClbP compared to no enzyme controls, indicating that the probe scaffold provides specific signal readout dependent on ClbP activity. Only a small amount of light (∼2% of WT) was observed from the ClbP S95A incubations compared to the blank (**Figure S2**). Detection of the hydrolyzed prodrug motif was found only in incubations with the wild type ClbP, indicating that the catalytic serine is necessary for proper activation of the probe (**Figure S3**).

**Figure 3.**
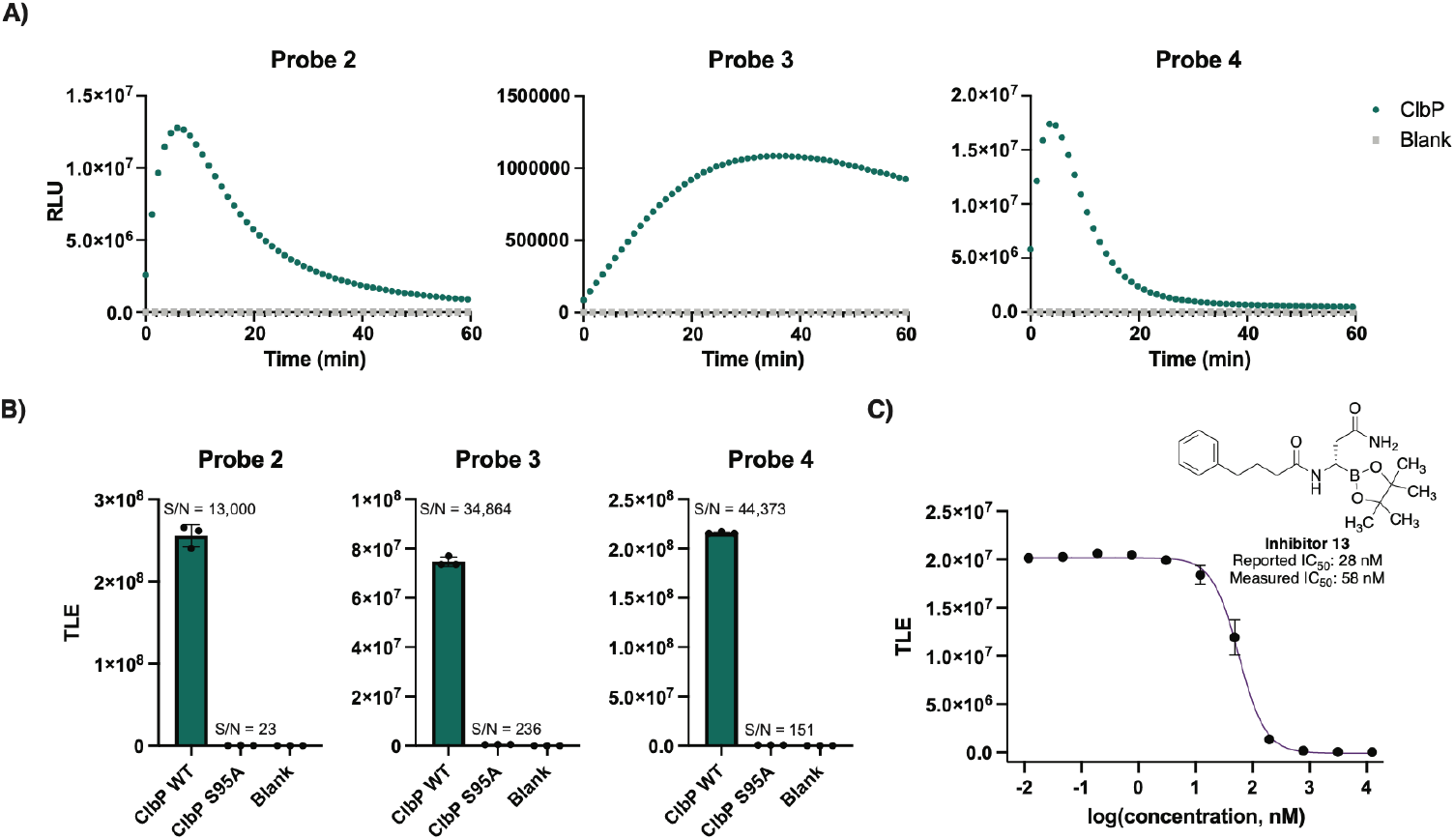
Chemiluminescent ClbP probes have increased sensitivity in vitro. A) Chemiluminescent kinetic profiles of probes 2-4 (10 µM) upon incubation with ClbP WT (13 nM) at room temperature. B) Comparison of total light emission of probes 2-4 after a 40 minute incubation with enzyme (13 nM for WT, 7.8 nM for S95A). C) Dose-response curve obtained by incubation of Probe 2 with Inhibitor 13 and light integration after 15 minutes. Error bars represent the ±SD of three independent measurements.

Light emissions from probes **2-4** yielded typical substrate processing curves, as well as signal-to-noise (S/N) ratios with at least five orders of magnitude for the samples containing WT ClbP (13 nM) above of blanks, and three orders of magnitude compared to ClbP S95A (7.8 nM) (**Figure 3B**). Notably, comparison of chemiluminescent probe **4** to the fluorescent probe **1** yielded a 770-fold increase of S/N at their respective maxima at equimolar concentrations (**Figure S4**). These results show that chemiluminescent probes produce a much stronger signal-to-noise readout when activated than a fluorescent probe. To evaluate the sensitivity of the chemiluminescent probes and previously described fluorescent probes in vitro, we measured the cumulative luminescence emitted by serial dilutions of ClbP after an hour of incubation with 10 µM of each probe. For probes **2-4**, we observed a limit of detection (LOD) at a concentration of 0.17 pM ClbP, a 625-fold improvement compared to that of the fluorescent probe (**Figure S5**). We also used previously reported inhibitor **13** to test the ability of chemiluminescence probes to report on inhibition of ClbP.^28^ We observed a dose-dependent reduction of emitted light that is inversely correlated with inhibitor concentration (**Figure 3C**), resulting in an IC_50_ value of 58 nM, close to that previously reported from fluorescent probe measurements (28 nM).^28^ These results show that the chemiluminescent probes **2**-**4** are able to report on ClbP activity in vitro. Lastly, we tested our chemiluminescent probes with a more distant homologue of ClbP, ZmaM, involved in the biosynthesis of Zwit-termicin which shares only 27% amino acid ID with ClbP. We observed that probe **2** reports on the activity of the WT protein, but not the catalytic serine mutant (S89A), with a slower rate than that observed for ClbP (**Figure S6**). This activity shows that the chemiluminescent probes can target enzymes other than ClbP from *E. coli* in vitro, with lesser identity to the canonical ClbP from *pks*^*+*^ *E. coli*.

### Testing of chemiluminescent ClbP probes in bacterial culture

Having confirmed the sensitivity of probes **2-4** for detecting activity of purified ClbP, we next sought to test their activity in live bacteria. ClbP is localized in the bacterial inner membrane, with its active site located at the interface of the periplasmic domain and the transmembrane helices of the protein.^24,37^ This localization requires that probes cross the bacterial outer membrane. To determine the effectiveness of probes **2-4** in bacteria, we incubated them individually with *E. coli* BW25113 heterologously expressing either the full *pks* gene cluster or the gene cluster with a Δ*clbP* deletion^26^ and measured luminescence. We observed an increase in luminescence after an hour only with the strain expressing the full *pks* gene cluster (**Figure 4A**). The Δ*clbP* background luminescence matched that of the blank controls (**Figure S7**). Integrated luminescence readings from probes **2**-**4** showed typical substrate consumption curves when incubated with *pks*^+^ bacteria (**Figure S8**). Of note, in incubations with similar amounts of bacteria, fluorescent probe **1** required a much longer time (> 2 hours) to produce signals detectable above background autofluorescence (**Figure S9**). Due to the delayed onset in luminescence, we performed RT-qPCR to probe changes in the transcripts of *clbP*. We did not observe differences between later time points relative to the starting time point (**Figure S10**). We hypothesize that a potential explanation is that the delay is caused by the time it takes for uptake to occur by the bacteria. These results indicate that *E. coli* cells activate probes chemiluminescent probes **2-4** more rapidly, and that the activation depends on the presence of catalytically active ClbP.

**Figure 4.**
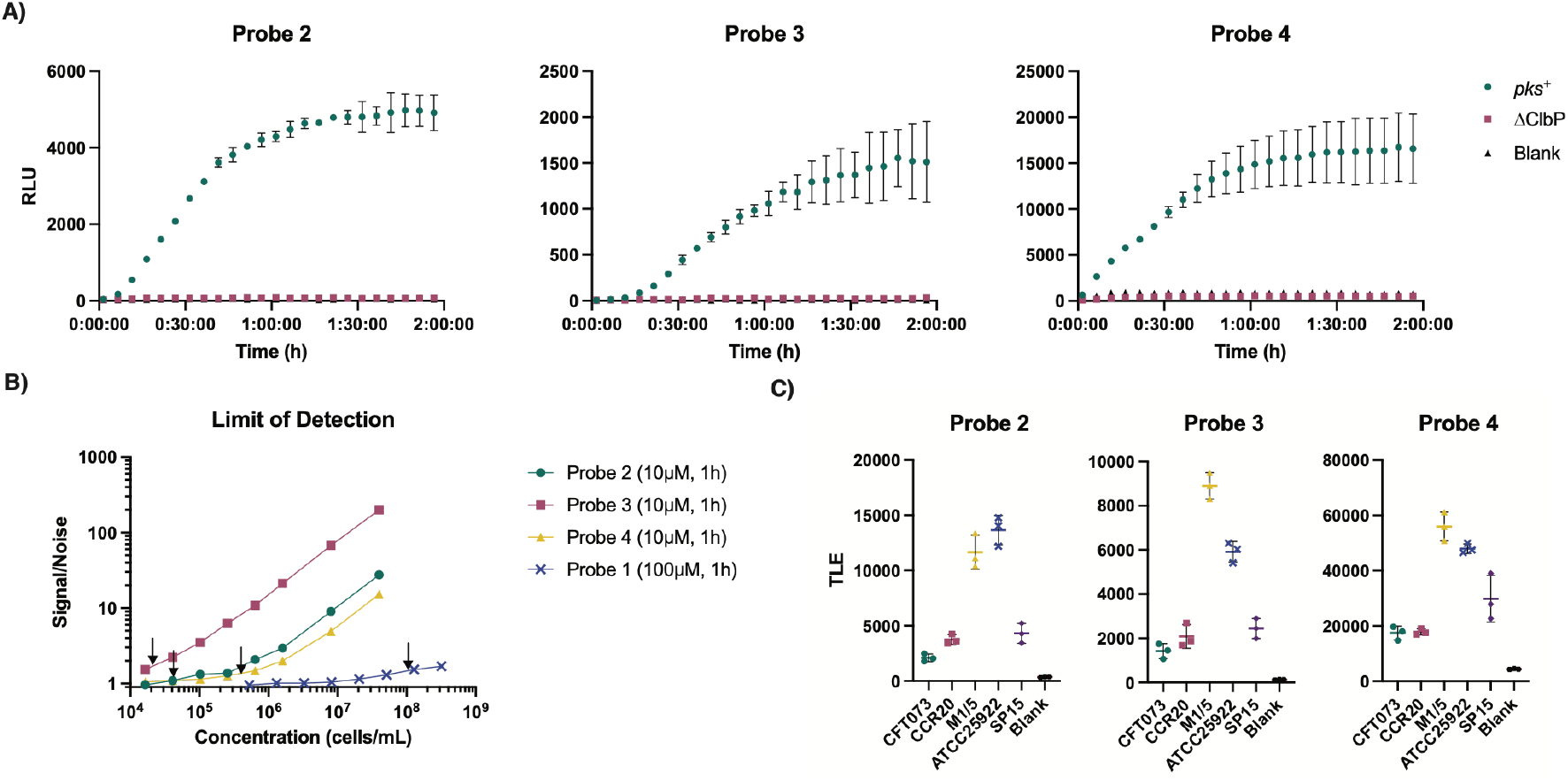
Chemiluminescent probes detect ClbP activity in *pks^+^ E. coli* cultures. A) Chemiluminescent kinetic profiles of probes 2-4 (10 µM) upon incubation with *E. coli* BW25113 heterologously expressing the *pks* gene cluster or a *ΔclbP* knockout in PBS. B) Determination of the limit of detection for a native *pks* encoder (*E. coli* Nissle 1917) with chemiluminescent probes 2-4 (10 µM) and fluorescent probe 1 (100 µM) after an hour at 37 ºC, with LOD values denoted by black arrows in each trace. C) Incubation of five *pks^+^ E. coli* isolates with probes 2-4 (10 µM) and light integration after one hour. Error bars represent the ±SD of three independent measurements.

To investigate the ability of the ClbP probes to detect lower numbers of bacteria in a more relevant setting, we used the native *pks*^+^ strain *E. coli* Nissle 1917. This strain has been used extensively in the study of colibactin^38–40^ and has the *pks* gene cluster under the control of its native regulators. We incubated serially diluted bacterial cultures with probes **2-4** (10 µM) and quantified the S/N ratio over time. We observed a marked increase in S/N between 30 minutes and 60 minutes, but no apparent benefit in averaging over 90 or 120 minutes (**Figure S11**). Therefore, we used 60-minute integrations for subsequent experiments. LODs for each probe were calculated by taking luminescence points with S/N < 15 and performing simple linear regressions to estimate the number of bacterial cells, which is equal to three times the standard deviation of a sample without cells; following previously reported methods.^41,42^ (**Figure S12**). These best fit lines showed a linear dependency between number of cells and cumulative light emission. Thus, probes **2-4** are quantitative for detection of *pks*^+^ bacteria. LODs were found to be 4.47×10^4^, 1.85×10^4^ and 3.07×10^5^ CFU/mL for probes **2-4**, respectively (**Figure 4B**). We performed a similar analysis for fluorescent probe **1**, in which we observed an extrapolated LOD of 9.75×10^7^ CFU/mL at 100 µM after an hour (**Figures 4B, S13-14**). This is >1000-fold less sensitive than probe **3** at 10 µM. These data clearly show that chemiluminescent ClbP probes have increased sensitivity over a fluorescent counterpart in bacterial cultures, even at lower probe concentrations.

We next examined the ClbP probes’ activity toward other native *pks*^+^ *E. coli*, with a 100% identical ClbP to *E. coli* Nissle 1917, including pathogenic *E. coli* isolates (CFT073, CCR20 and SP15) and *E. coli* isolates from healthy volunteers (M1/5 and ATCC 25922) by measuring the luminescence produced after a 1-hour incubation with probe (**Figure 4C**). All three chemiluminescent probes reliably produced luminescence above background levels in the presence of all five strains, whereas fluorescent probe **1** did not have detectable activity for every strain after an hour (**Figure S15**). Finally, we incubated *E. coli* Nissle 1917 with probes **2-4** in the presence of ClbP inhibitor **13** (10 µM) to test whether the probes can report on inhibition of ClbP in bacterial cultures. We observed a complete reduction of luminescence when bacterial cultures were incubated with probe and inhibitor **13** (**Figure S16**). This is consistent with previous results obtained with fluorescent probe **1**^43^. Overall, probes **2-4** can report on the activity of native *pks*^+^ *E. coli*, including detecting their inhibition by small molecules. To survey a more diverse group of bacteria for their activity towards the new chemiluminescent probes, we looked at other encoders of the *pks* cluster. *Frischella perrara* (52% amino acid ID to *E. coli* ClbP) and *Erwinia oleae* (87% amino acid ID to *E. coli* ClbP) are two known producers of colibactin^44,45^, whilst *Pseudovibrio denitrificans*, (29% amino acid ID to *E. coli* ClbP) carries a more dissimilar *pks* gene cluster that has not been functionally characterized. Multiple sequence alignment of these ClbP show complete conservation of the active site triad and maintenance of the auxiliary amino acids that interact with the asparagine sidechain and lipid tail of the prodrug motif, highlighting that our probes would likely be able to interact with the proteins (**Figure S17**). To validate the use of our probes with strains other than *E. coli* we will use *Klebsiella pneumoniae* (the second most abundant Enterobacteriaceae in the human gut^46^) strains WGLW3 (*pks*^*+*^, 100% ID to *E. coli* ClbP) and WGLW5 (*pks*^*–*^), a member of the Erwiniaceae family, *Erwinia oleae* (strain DAPP-PG-531, 87% ID) and an alphaproteobacterium, *Pseudovibrio denitrificans* (strain JCM 12308, 29% ID) which bears the least identical ClbP homologue known in a *pks* cluster. We observed that all probes were able to be processed by the *pks*^*+*^ organisms only (**Figure S18)**. Furthermore, we also observed that this activity is ClbP related, as the ClbP inhibitor **13** prevents the luminescence for all the different bacteria (**Figure S19**). Testing of chemiluminescent probes in biologically complex samples

In light of their high sensitivity and selectivity, we asked whether our chemiluminescent ClbP probes could detect *pks*^+^ gut bacteria directly in a complex microbial community. We envisioned that they might enable a simplified procedure for rapid detection of *pks*^+^ organisms that would not require intensive anaerobic culturing (**Figure 5A**). Previously, other chemiluminescent assays have been used as readouts to detect the presence of occult blood in stool (through hemoglobin-catalyzed oxidation of luminol)^47^ and to detect *K. pneumoniae* spiked into fecal samples (using bioluminescent phages).^48^ Probes **2-4** differ from these examples as they rely on detecting the activity of a gut microbial enzyme.

**Figure 5.**
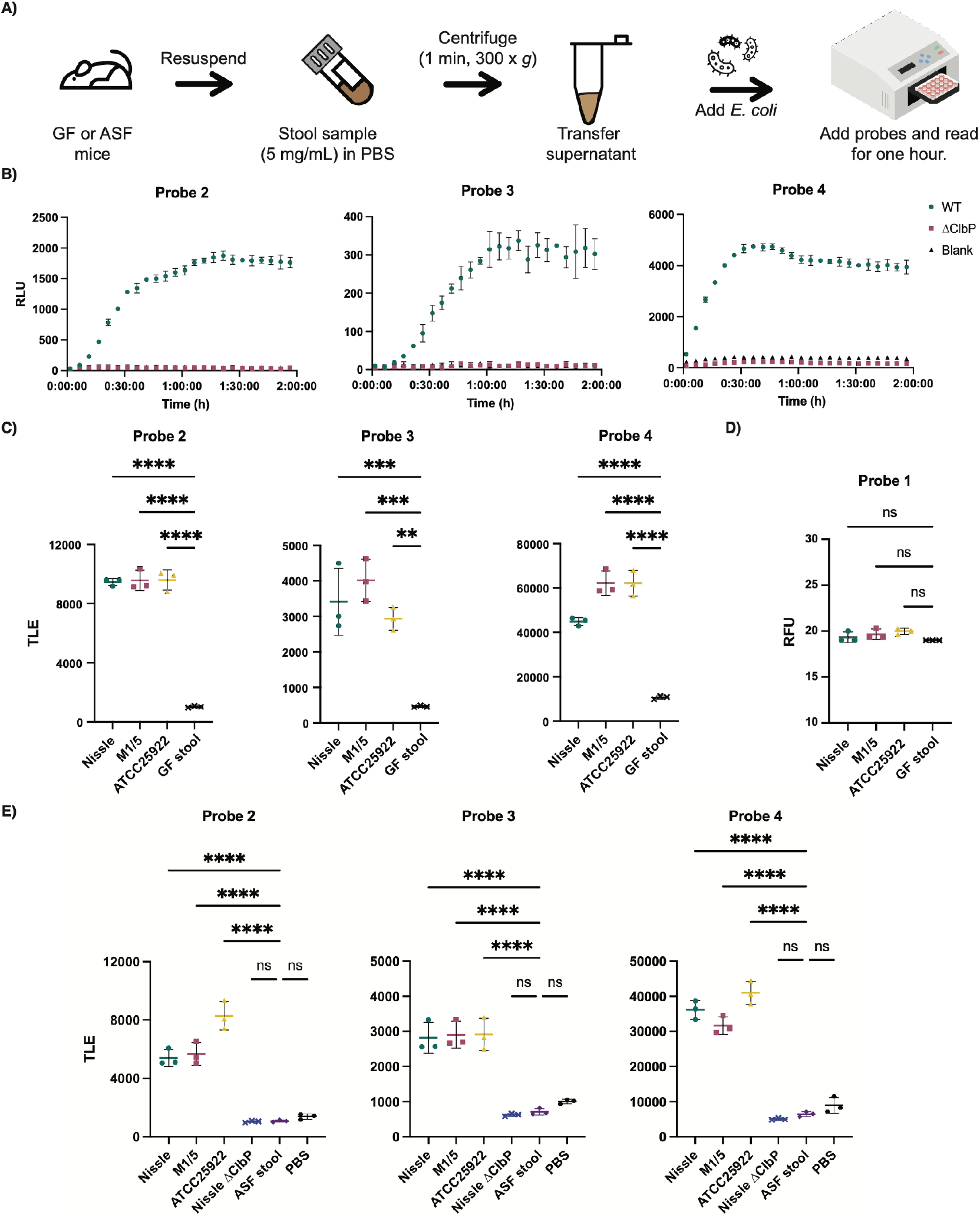
Chemiluminescent probes allow for facile detection of *pks^+^ E. coli* in biological samples. A) Workflow for processing stool samples in PBS (pH 7.4) for chemiluminescent assays. B) Chemiluminescent kinetic profiles of 2-4 (10 µM) upon incubation with E. coli BW25113 bearing the full *pks* gene cluster or a *ΔclbP* knockout in a germ-free (GF) stool resuspension in PBS C) Detection of natural pks isolates with chemiluminescent probes 2-4 (at 25 µM) in GF stool resuspensions and total light integration after one hour. D) Detection of natural pks isolates with fluorescent probe 1 (at 100 µM) in GF stool resuspensions and fluorescence after one hour. E) Detection of natural pks isolates with chemiluminescent probes 2-4 (at 25 µM) in ASF stool resuspensions and total light integration after one hour. Error bars represent the ±SD of three independent measurements. ****P<0.0001; ***P<0.001; **P<0.01; Not significant (ns) *P*>0.05 using one-way ANOVA and Dunnett’s multiple comparison test.

Detection of *pks*^+^ gut bacteria in stool requires the chemiluminescent probes to remain stable in this complex sample type. To assess probe stability, we initially incubated probes with PBS suspensions of stool from germ-free mice. No significant differences in luminescence were observed between control solutions and stool suspensions up to 5 mg/mL of stool (**Figure S20**). This indicates that probes are stable to the endogenous contents of mouse stool. To ensure that light emission was not impeded by the fecal matter, we added either recombinant WT or S95A ClbP to germ-free stool resuspensions (10 nM) and observed light production only for samples containing WT ClbP (**Figure S21**).

To optimize sample preparation, we removed particulates in germ-free stool suspensions using filters of different sizes. No significant differences in luminescence were observed between the filtered and unfiltered samples (**Figure S22**), indicating that filtering was not necessary for sample preparation. Similarly, adding *E. coli* BW25113 to germ-free fecal samples resulted in robust light production after an hour only with the strain expressing WT ClbP (**Figure 5B** and **Figure S23**). Using this same assay format, fluorescent probe **1** did not show an increase in fluorescence over background (**Figure S24**).

To ensure the sensitivity of the assay, we tested probes **2**-**4** in stool suspensions from germ-free mice and added anaerobically-grown native *pks*^+^ gut bacteria (Nissle 1917, M1/5, ATCC 25992) at levels reflecting those at which *E. coli* is normally found in humans (∼ 10^7^ to 10^9^ CFU per gram of wet stool).^49^ We found robust activation in the samples with bacteria compared to that of the background within an hour (**Figure 5C**). In contrast, we did not observe a noticeable increase in fluorescence over background for probe **1** in this assay format within this timeframe (**Figure 5D**). Furthermore, we applied the same assay to stool obtained from germ-free mice colonized with Altered Schaedler Flora (ASF).^50^ We observed a significant increase of luminescence over blank only with the addition of *pks*^*+*^ bacteria (**Figure 5E**). Therefore, these results provide a proof-of-concept that chemiluminescent probes **2-4** can be applied to detect the presence of native *pks*^+^ organisms in complex samples.

## Conclusions

In summary, we have developed chemiluminescent probes for the facile detection of the essential colibactin biosynthetic enzyme ClbP. Informed by previous studies of ClbP’s activity and earlier probe development efforts,^26,27^ these second generation probes are more sensitive (up to >1000 fold) than prior fluorescent ClbP probes. This sensitivity allows for the detection of ClbP activity in more complex biological samples, including stool. Though probe **3** bears a methyl ester, which is known to aid outer membrane permeability^51^, probes **2** and **4** are able to enter the periplasm as well. We note that these probes are also processed differently in vitro, with **2** and **4** being metabolized by the enzyme with a faster T_max_ than probe **3**. Future studies will focus on improving on probe **4**, as it is the brightest of these probes. The increased sensitivity of our chemiluminescent probes also greatly reduces assay times (<1 hour) compared to previously reported fluorescent probes.

Though there are many potential explanations for colibactin production by bacteria, current hypotheses focus on its role in competition against other microbes and host as collateral damage. Recent examples include its antagonistic activity against *Phocaeicola vulgatus* (a commensal organism), *Vibrio cholerae (*a pathogen) and *Bacteroides fragilis* in the human microbiome^52,53^, and *Serratia marcescens* (a pathogen) in the honey-bee gut microbiome^54^.The accumulating evidence that *pks*^*+*^ bacteria may contribute to tumorigenesis in humans highlights a potential need to detect the presence of these organisms in patients for cancer prevention and/or diagnosis. Though our probes can report activity in a model stool resuspension experiment, further testing in complex fecal samples containing pks+ organisms is required to verify their utility as a point-of-care diagnostic. An ideal test for the presence of *pks*^*+*^ bacteria necessitates both sensitivity and specificity, as well as providing outcomes in a timely manner. Though prior fluorescent probes are specific to *pks*^*+*^ organisms, they lack the sensitivity and speed associated with chemiluminescent probes **2**-**4**. Furthermore, the chemiluminescent assay has been miniaturized (to 384-wells plates) which could allow for faster parallel testing of clinical samples for *pks*^*+*^ bacteria.

Another advantage of these chemiluminescent probes is that they can report quickly on the inhibition of ClbP activity (< 1 hour), a benefit that can be exploited for high-throughput screening of ClbP inhibitor candidates in bacterial samples.^29^ This would increase the throughput with which small molecule libraries can be screened, compared to that of fluorescent probes or mass spectrometry-based methods, which required up to 72 hours to produce a readable signal.^29^

The development of next-generation ClbP probes adds to the growing evidence that chemiluminescent probes targeting bacterial enzymes are exquisitely selective and sensitive.^30,33,34^ Future probe optimization could include incorporation of brighter luminophores^41^ and production of red-shifted light for *in vivo* spatially-resolved monitoring of ClbP activity in tissues^55^. Altogether, our work highlights the importance of developing sensitive and specific chemical probes to track health-relevant gut bacterial enzymatic activities in complex samples and will facilitate future efforts to elucidate the importance of specific bacterial activities in human health and diseases.

## ASSOCIATED CONTENT

### Supporting Information

The Supporting Information is available free of charge on the ACS Publications website.

Experimental details, supplemental figures, synthetic schemes, and compound characterization (PDF)

## AUTHOR INFORMATION

### Present Addresses

†If an author’s address is different than the one given in the affiliation line, this information may be included here.

### Author Contributions

The manuscript was written through contributions of all authors. / All authors have given approval to the final version of the manuscript.

## ACKNOWLEDGMENT

We thank C. Liu and X. Dong for insightful discussion and critical reading of the manuscript, G. Zavala and R. Gaudet for providing recombinant ZmaM, and W. Garrett for sharing samples of mouse fecal pellets. The following reagents were obtained through BEI Resources, NIAID, NIH as part of the Human Microbiome Project: *Klebsiella pneumoniae subsp. pneumoniae*, Strain WGLW3, HM-748; and Strain WGLW5, HM-749. Elements of selected figures were created with Biorender.com. This article is subject to HHMI’s Open Access to Publications policy. HHMI lab heads have previously granted a nonexclusive CC BY 4.0 license to the public and a sublicensable license to HHMI in their research articles. Pursuant to those licenses, the author-accepted manuscript of this article can be made freely available under a CC BY 4.0 license immediately upon publication.

## Insert Table of Contents artwork here

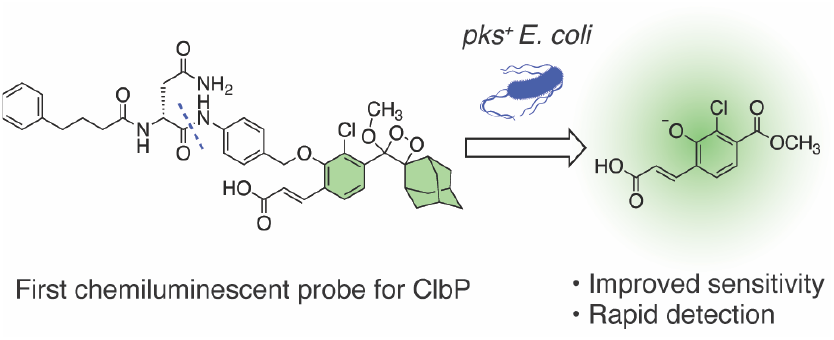

