## Supplementary information for "Chemiluminescent probes allow for the rapid identification of colibactin-producing bacteria"

Measurements of probe activity in stool samples containing native *pks*[+] *E. coli* isolates ... 20

#### General materials and methods

All chemicals were purchased from Sigma Aldrich (St. Louis, MO) unless otherwise noted. Luria-Bertani (LB) media was purchased from Alfa-Aesar (Ward Hill, MA). Casamino acids was purchased from VWR (West Chester, PA). PBS (1X, pH 7.4) was purchased from Thermo Fisher Scientific (Waltham, MA). Trace Mineral Supplement and Vitamin Supplement were purchased from ATCC (Manassas, VA). Nutrient Broth (NB), Tryptic Soy Broth (TSB) and Marine Broth (MB) were purchased from BD Difco (Franklin Lakes, NJ). Solvents used for LC–MS were B&J Brand high-purity solvents (Honeywell, Charlotte, NC).

Strain of *E. coli* Nissle 1917 and *E. coli* Nissle 1917 ΔClbP was received from the Müller lab (Helmholtz Institute of Pharmaceutical Research, Germany), *E. coli* CCR20, *E. coli* BW25113 pBeloBAC11-*pks* and *E. coli* BW25113 pBeloBAC11-*pksΔclbP* was received from the Bonnet lab (Laboratoire de Bactériologie Clinique, France), *E. coli* CFT073, SP15 and M1/5 were received from the Dobrindt lab (University of Münster, Germany), and *E. coli* ATCC 25922 was purchased from ATCC (Manassas, VA). *Klebsiella pneumoniae subsp. pneumoniae* strain WGLW3 and WGLW5 were obtained from BEI resources (Manassas, VA), *Erwinia oleae* DAPP-PG-531 and *Pseudovibrio denitrificans* JCM 12308 were obtained from DSMZ (Braunschweig, Germany).

Optical densities of bacterial cultures were determined by measuring their absorbance at 600 nm with a DU 730 UV/Vis spectrophotometer (Beckman Coulter, Indianapolis, IN). Fluorescent measurements were performed on a BioTek Synergy HTX microplate reader (Winooski, VT) and chemiluminescent measurements were performed on a Molecular Devices Spectramax iD3 (San Jose, CA) or a BioTek Synergy Neo2 microplate reader (Winooski, VT). Plates utilized were Corning 3570 or 3575 (Corning, NY) for microplate reader assays.

Anaerobic bacterial growth was performed in an anaerobic chamber (Coy Laboratories, Grass Lake, MI) under a 95% $N_2$ and 5% $H_2$ atmosphere. Solutions (<50 mL), media, and plastics were made anaerobic by bringing them into the anaerobic chamber and equilibrating for at least 48 hours before usage.

#### Synthetic methods

All reactions requiring anhydrous conditions were performed under an argon atmosphere. All reactions were carried out at room temperature unless stated otherwise. Chemicals and solvents were either A.R. grade or purified by standard techniques. Thin-layer chromatography (TLC): silica gel plates Merck 60 F254, compounds were visualized by irradiation with UV light. Column chromatography (FC): silica gel Merck 60 (particle size 0.040-0.063 mm), eluent given in parentheses. Reverse-phase high-pressure liquid chromatography (RP-HPLC): C18 5 μm, 250 x 4.6 mm, eluent given in parentheses. Preparative RP-HPLC: C18 5 μM, 250 x 21mm, eluent given in parentheses. $^{1}$H NMR spectra were measured using a Bruker Avance operated at 400 MHz. $^{13}$C NMR spectra were measured using a Bruker Avance operated at 101 MHz. Chemical shifts were reported in ppm on the δ scale relative to a residual solvent ($CDCl_3$: δ = 7.26 for $^{1}$H NMR and 77.16 for $^{13}$C NMR, MeOH-$d_4$: δ = 3.31 for $^{1}$H NMR and 49.00 for $^{13}$C NMR, DMSO-$d_6$: δ = 2.50 for $^{1}$H-NMR and 39.52 for $^{13}$C NMR, Acetic Acid-d4: δ = 2.04 and 11.65 for $^{1}$H-NMR and 20.0 and 178.99 for $^{13}$C NMR). Mass spectra were measured on Waters Xevo SQD2 and Waters Xevo G2-XS Q-TOF (Milford, MA). All general reagents, including salts and solvents, were purchased from Sigma-Aldrich. Light irradiation for photochemical reactions: LED PAR38 lamp (19W, 3000K).

#### Abbreviations

**ACN**- Acetonitrile, **DCM**- Dichloromethane, **DMF**- *N,N'*–Dimethylformamide, **EEDQ**- *N*-Ethoxycarbonyl-2-ethoxy-1,2-dihydroquinoline, **$Et_3N$**– Triethylamine, **$Et_2O$**- Diethyl ether, **EtOAc**- Ethyl acetate, **Fmoc**- Fluorenylmethyloxycarbonyl, **Hex**- Hexanes, **MB**- Methylene blue, **MeOH**- Methanol, **TFA**- Trifluoroacetic acid, **THF**- Tetrahydrofuran **TMSCl**- Trimethylsilyl chloride, **CFU**– Colony forming units, **$OD_{600}$**– Optical Density at 600 nm, **PBS**- Phosphate buffered saline (pH 7.4).

#### Synthesis and characterization

Compound **5**

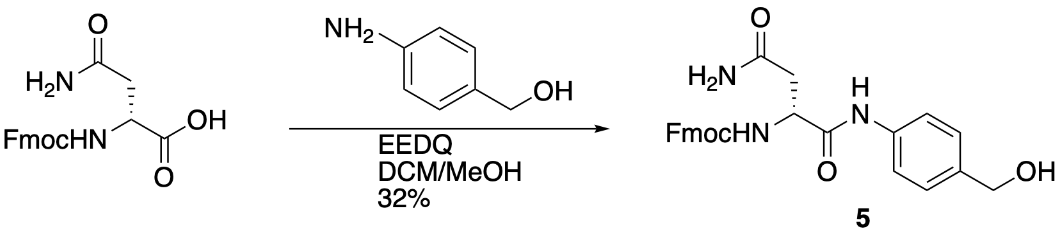

Fmoc-D-asparagine (1.2 g, 3.39 mmol, 1 eq.) and 4-aminobenzyl alcohol (0.5 g, 4.06 mmol, 1.2 eq.) were partially dissolved in 12 mL of 1:1 DCM/MeOH. EEDQ (1.26 g, 5.08 mmol, 1.5 eq.) was added, and the solution was stirred overnight. The reaction was monitored by TLC. Upon completion, $Et_2O$ was added to the reaction solution, and the crude product was isolated by vacuum filtration. The product was purified by silica gel column chromatography (95:5 EtOAc: MeOH) to afford compound **5** as a brown solid (495 mg, 32% yield).

**$^1$H NMR (400 MHz, DMSO)** δ 9.98 (s, 1H), 7.89 (d, *J* = 7.5 Hz, 2H), 7.73 (d, *J* = 7.4 Hz, 2H), 7.65 – 7.54 (m, 3H), 7.42 (t, *J* = 7.4 Hz, 2H), 7.36 – 7.29 (m, 3H), 7.23 (d, *J* = 8.4 Hz, 2H), 6.94 (s, 1H), 5.09 (t, *J* = 5.7 Hz, 1H), 4.52 – 4.46 (m, 1H), 4.43 (d, *J* = 5.6 Hz, 2H), 4.34 – 4.16 (m, 4H).

**$^{13}$C NMR (101 MHz, DMSO)** δ 171.14, 170.07, 155.80, 143.83, 140.72, 137.62, 137.44, 127.66, 127.11, 126.83, 125.33, 120.12, 119.07, 65.75, 62.60, 52.38, 46.63, 37.32.

**MS (ES+):** m/z calc. for $C_{26}H_{25}N_3O_5$: 459.2; found: 482.2 $[M+Na]^+$.

<u>Compound **6**</u>

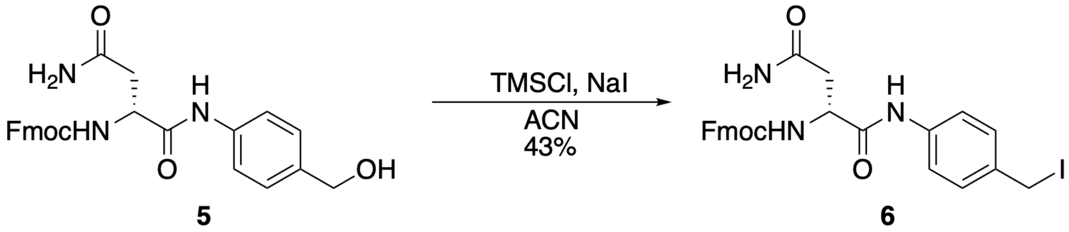

Compound **5** (200 mg, 0.435 mmol, 1 eq.) was partially dissolved in 2 mL of ACN. NaI (196 mg, 1.31 mmol, 3 eq.) and TMSCl (166 µL, 1.31 mmol, 3 eq.) were added, and the solution was stirred at room temperature. The reaction was monitored by TLC. Once most of the starting material had been consumed, the reaction mixture was diluted with EtOAc and washed with saturated $NH_4Cl$ and $Na_2S_2O_3$. The organic phase was dried over $Na_2SO_4$, and the solvent was evaporated under reduced pressure. The product was purified by silica gel column chromatography (30:70 Hex: EtOAc) to afford compound **6** as a yellow solid (106 mg, 43% yield).

**$^1$H NMR (400 MHz, Acetic acid)** δ 7.79 (d, *J* = 6.9 Hz, 2H), 7.68 – 7.53 (m, 4H), 7.43 – 7.14 (m, 7H), 5.05 – 4.62 (m, 2H), 4.57 – 4.20 (m, 5H), 3.01 – 2.83 (m, 2H).

**$^{13}$C NMR (101 MHz, Acetic acid)** δ 175.38, 170.43, 157.08, 143.68, 141.20, 136.29, 129.14, 127.60, 126.98, 125.00, 120.66, 119.81, 115.67, 67.30, 52.58, 46.91, 29.47, 13.30.

**MS (ES+):** m/z calc. for $C_{26}H_{24}IN_3O_4$: 569.1; found: 592.6 $[M+Na]^+$.

<u>Compound **8**</u>

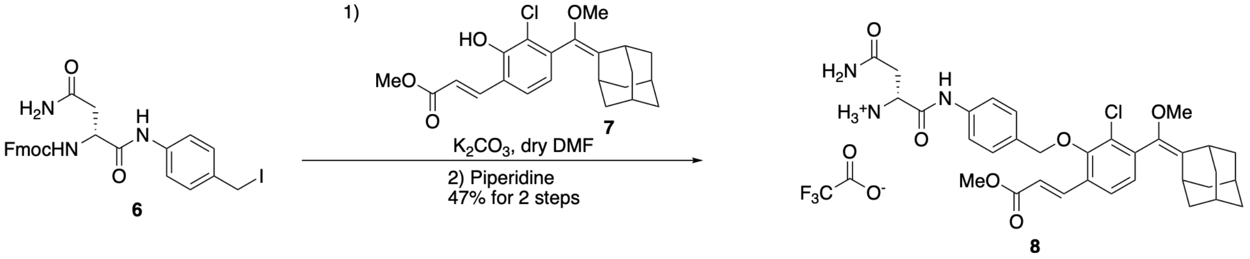

Compound **6** (54 mg, 0.095 mmol, 1 eq.), compound **7**[1] (37 mg, 0.095 mmol, 1 eq.), and $K_2CO_3$ (26 mg, 0.190 mmol, 2 eq.) were dissolved in dry DMF. The reaction was monitored by RP-HPLC (70-100% ACN in water, 0.1% TFA). Upon completion, 200 µL of piperidine was added to the solution. The deprotection of the Fmoc was monitored by RP-HPLC (70-100% ACN in water, 0.1% TFA). Upon completion, the piperidine and the DMF were evaporated under reduced pressure. The product was purified by preparative RP-HPLC (50-100% ACN in water, 0.1% TFA) to afford compound **8** as a white solid (32 mg, 47% yield).

**$^1$H NMR (400 MHz, MeOD)** δ 7.85 (d, *J* = 16.2 Hz, 1H), 7.62 – 7.56 (m, 3H), 7.42 – 7.36 (m, 2H), 7.11 (d, *J* = 8.0 Hz, 1H), 6.49 (d, *J* = 16.2 Hz, 1H), 5.00 (d, *J* = 6.1 Hz, 2H), 4.35 (dd, *J* = 8.8, 4.6 Hz, 1H), 3.79 (s, 3H), 3.30 (s, 3H), 3.25 (s, 1H), 3.05 – 2.98 (m, 1H), 2.91 – 2.83 (m, 1H), 2.47 (s, 1H), 2.33 (d, *J* = 14.4 Hz, 1H), 2.08 – 1.58 (m, 12H).

**$^{13}$C NMR (101 MHz, MeOD)** δ 173.24, 168.74, 167.59, 154.83, 140.99, 139.96, 139.48, 139.30, 133.64, 133.05, 131.26, 131.06, 130.77, 129.13, 126.40, 120.85, 120.79, 76.77, 57.42, 52.33, 51.99, 48.36, 40.36, 38.40, 38.11, 34.75, 34.40, 34.24, 31.05, 28.88, 28.56.

**MS (ES+):** m/z calc. for $C_{33}H_{38}ClN_3O_6$: 607.2; found: 608.8 $[M+H]^+$.

<u>Compound **9**</u>

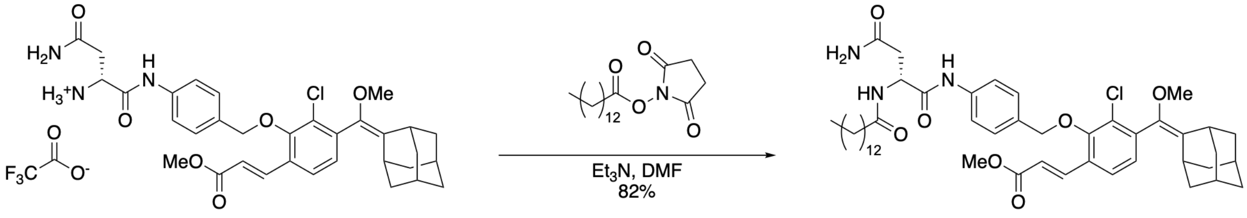

Compound **8** (32 mg, 0.044 mmol, 1 eq.), 2,5-dioxopyrrolidin-1-yl tetradecanoate[2] (25 mg, 0.078 mmol, 2 eq.), and $Et_3N$ (16.2 µL, 0.116 mmol, 3 eq.) were dissolved in DMF. The reaction was monitored by RP-HPLC (70-100% ACN in water, 0.1% TFA). Upon completion, the reaction mixture was diluted with EtOAc and washed with brine. The organic phase was dried over $Na_2SO_4$, and the solvent was evaporated under reduced pressure. The product was purified by silica gel column chromatography (100% EtOAc) to afford compound **9** as a colorless gel (26 mg, 82% yield).

**$^{1}$H NMR (400 MHz, $CDCl_3$)** δ 9.59 (s, 1H), 7.89 (d, *J* = 16.2 Hz, 1H), 7.61 (d, *J* = 6.8 Hz, 1H), 7.54 (d, *J* = 8.4 Hz, 2H), 7.42 (t, *J* = 7.9 Hz, 3H), 7.07 (d, *J* = 8.0 Hz, 1H), 6.49 (s, 1H), 6.42 (d, *J* = 16.2 Hz, 1H), 5.83 (s, 1H), 4.98 – 4.87 (m, 3H), 3.80 (s, 3H), 3.33 (s, 3H), 3.28 (s, 1H), 2.93 (dd, *J* = 15.5, 3.5 Hz, 1H), 2.64 (dd, *J* = 15.5, 6.9 Hz, 1H), 2.30 (t, *J* = 7.6 Hz, 2H), 2.07 (s, 1H), 1.99 – 1.57 (m, 12H), 1.31 – 1.19 (m, 22H), 0.88 (t, *J* = 6.8 Hz, 3H).

**$^{13}$C NMR (101 MHz, $CDCl_3$)** δ 174.29, 174.17, 169.14, 167.14, 153.64, 139.41, 138.89, 138.15, 138.04, 132.40, 131.95, 129.83, 129.73, 127.77, 125.07, 119.84, 75.71, 57.24, 51.83, 50.59, 39.19, 39.04, 38.61, 37.06, 36.54, 36.27, 32.94, 31.92, 31.63, 29.65, 29.50, 29.35, 29.27, 28.35, 28.20, 25.58, 22.69, 14.12.

**MS (ES+):** m/z calc. for $C_{47}H_{64}ClN_3O_7$: 817.4; found: 841.1 $[M+Na]^+$.

Compound **11**

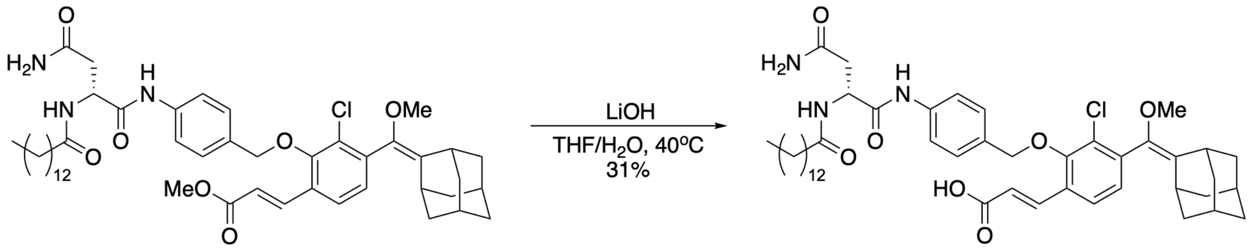

9 11

Compound **9** (26 mg, 0.032 mmol, 1 eq.) was dissolved in 1 mL of 4:1 THF/$H_2O$. LiOH (7.6 mg, 0.318 mmol, 10 eq.) was added and the reaction was heated to 40°C. The reaction was monitored by RP-HPLC (90-100% ACN in water, 0.1% TFA). Upon completion, the solvent was evaporated under reduced pressure. The product was purified by preparative RP-HPLC (90-100% ACN in water, 0.1% TFA) to afford compound **11** as a colorless gel (8 mg, 31% yield).

**$^1$H NMR (400 MHz, $CDCl_3$)** δ 9.65 – 9.52 (m, 1H), 7.75 (s, 1H), 7.64 (d, *J* = 15.9 Hz, 1H), 7.53 – 7.31 (m, 4H), 7.24 (d, *J* = 7.9 Hz, 2H), 7.04 (d, *J* = 7.9 Hz, 1H), 6.83 (s, 1H), 6.17 (d, *J* = 16.1 Hz, 1H), 5.07 (s, 1H), 4.98 – 4.89 (m, 2H), 3.33 (s, 3H), 3.28 (s, 1H), 2.81 (s, 2H), 2.29 – 2.19 (m, 3H), 2.08 (s, 1H), 1.99 – 1.51 (m, 11H), 1.24 – 1.16 (m, 22H), 0.85 (t, *J* = 6.1 Hz, 3H).

**$^{13}$C NMR (101 MHz, $CDCl_3$)** δ 175.23, 169.94, 153.87, 140.34, 139.58, 138.44, 138.08, 132.66, 132.20, 130.34, 130.16, 129.80, 128.30, 127.91, 124.85, 124.11, 120.48, 119.30, 76.43, 57.40, 51.16, 39.37, 38.80, 37.19, 36.51, 33.12, 32.05, 31.72, 31.57, 30.00, 29.83, 29.66, 29.49, 29.39, 28.48, 25.73, 22.82, 14.25.

**MS (ES+):** m/z calc. for $C_{46}H_{62}ClN_3O_7$: 803.4; found: 827.0 $[M+Na]^+$.

Probe **2**

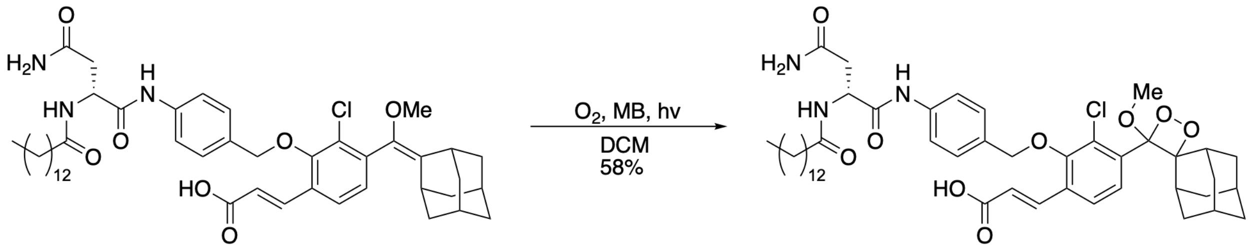

11 Probe 2

Compound **11** (5 mg, 0.006 mmol) and a catalytic amount of methylene blue were dissolved in 10 mL of DCM. Then, oxygen was bubbled through the solution while irradiating with yellow light. The reaction was monitored by RP-HPLC (90-100% ACN in water, 0.1% TFA). Upon completion, the solvent was evaporated under reduced pressure. The crude product was passed through a silica gel column (100 EtOAc with a few drops of AcOH) to afford **Probe 2** as a colorless gel (3 mg, 58% yield).

**$^1$H NMR (400 MHz, DMSO)** δ 10.01 (s, 1H), 8.09 (d, *J* = 7.7 Hz, 1H), 7.91 (d, *J* = 8.4 Hz, 1H), 7.77 – 7.69 (m, 2H), 7.60 (d, *J* = 8.5 Hz, 2H), 7.35 (d, *J* = 8.4 Hz, 2H), 7.30 (s, 1H), 6.87 (s, 1H), 6.62 (d, *J* = 16.1 Hz, 1H), 4.92 – 4.80 (m, 2H), 4.66 (dd, *J* = 14.0, 7.2 Hz, 1H), 3.09 (s, 3H), 2.86 (s, 1H), 2.22 (d, *J* = 11.6 Hz, 1H), 2.09 (t, *J* = 7.3 Hz, 2H), 2.01 – 1.95 (m, 1H), 1.89 (s, 1H), 1.72 – 1.40 (m, 12H), 1.24 – 1.14 (m, 22H), 0.82 (t, *J* = 6.5 Hz, 3H).

**$^{13}$C NMR (101 MHz, DMSO)** δ 172.78, 171.63, 170.64, 167.85, 153.92, 139.80, 134.20, 132.07, 130.76, 129.76, 128.79, 127.27, 126.44, 119.50, 111.66, 95.86, 75.87, 51.09, 49.84, 37.56, 36.34, 35.58, 33.74, 33.51, 32.30, 32.13, 31.76, 31.61, 31.33, 30.28, 29.53, 29.49, 29.32, 29.18, 29.05, 25.97, 25.67, 22.55, 14.41.

**MS (ES-):** m/z calc. for $C_{46}H_{62}ClN_3O_9$: 835.4; found: 835.0 $[M-H]^-$.

<u>Compound **10**</u>

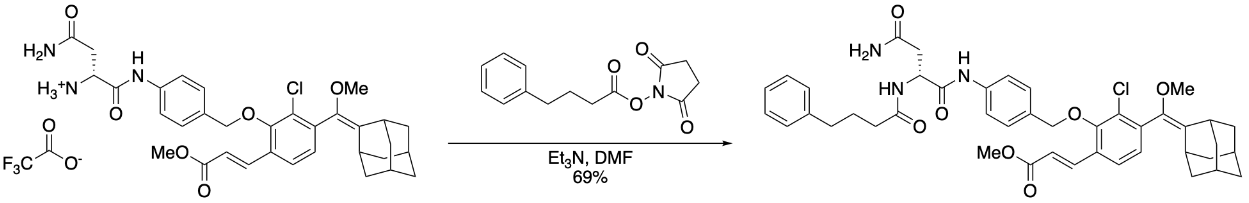

**8** **10**

Compound **8** (32 mg, 0.044 mmol, 1 eq.), 2,5-dioxopyrrolidin-1-yl 4-phenylbutanoate[3] (23 mg, 0.089 mmol, 2 eq.), and $Et_3N$ (18.5 µL, 0.133 mmol, 3 eq.) were dissolved in DMF. The reaction was monitored by RP-HPLC (70-100% ACN in water, 0.1% TFA). Upon completion, the $Et_3N$ and DMF were evaporated under reduced pressure. The product was purified by preparative RP-HPLC (70-100% ACN in water, 0.1% TFA) to afford compound **10** as a white solid (23 mg, 70% yield).

**$^1$H NMR (400 MHz, CDCl$_3$)** δ 9.54 (s, 1H), 7.85 (d, *J* = 16.2 Hz, 1H), 7.54 – 7.48 (m, 3H), 7.38 (t, *J* = 8.0 Hz, 3H), 7.23 (d, *J* = 8.3 Hz, 2H), 7.19 – 7.11 (m, 3H), 7.04 (d, *J* = 8.0 Hz, 1H), 6.44 – 6.35 (m, 2H), 5.89 (s, 1H), 4.96 – 4.84 (m, 3H), 3.76 (s, 3H), 3.30 (s, 3H), 3.26 (s, 1H), 2.85 (dd, *J* = 15.5, 3.5 Hz, 1H), 2.66 – 2.56 (m, 4H), 2.27 (t, *J* = 7.5 Hz, 2H), 2.05 (s, 1H), 1.99 – 1.62 (m, 13H).

**$^{13}$C NMR (101 MHz, CDCl$_3$)** δ 174.35, 174.02, 169.16, 167.31, 153.76, 141.30, 139.54, 139.03, 138.30, 138.12, 132.56, 132.15, 129.95, 129.87, 128.59, 127.92, 126.20, 125.20, 120.01, 119.93, 75.84, 57.38, 51.96, 50.72, 39.32, 39.18, 38.75, 37.19, 36.62, 35.81, 35.29, 33.08, 29.84, 28.49, 28.34, 27.10.

**MS (ES+):** m/z calc. for $C_{43}H_{48}ClN_3O_7$: 753.3; found: 777.0 $[M+Na]^+$.

<u>Probe **3**</u>

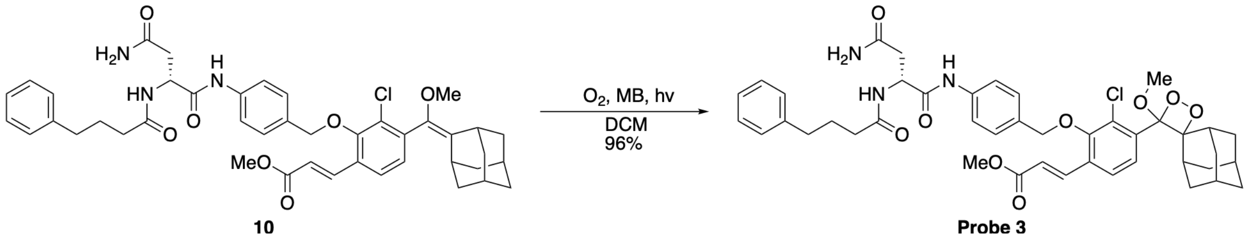

Compound **10** (13 mg, 0.017 mmol) and a catalytic amount of methylene blue were dissolved in 10 mL of DCM. Then, oxygen was bubbled through the solution while irradiating with yellow light. The reaction was monitored by RP-HPLC (70-100% ACN in water, 0.1% TFA). Upon completion, the solvent was evaporated under reduced pressure. The product was purified by preparative RP-HPLC (70-100% ACN in water, 0.1% TFA) to afford **Probe 3** as a white solid (13 mg, 96% yield).

**$^1$H NMR (400 MHz, CDCl$_3$)** δ 9.51 (s, 1H), 7.92 (d, *J* = 8.4 Hz, 1H), 7.85 (dd, *J* = 16.2, 1.7 Hz, 1H), 7.59 – 7.49 (m, 4H), 7.39 (dd, *J* = 8.4, 1.1 Hz, 2H), 7.31 – 7.25 (m, 2H), 7.22 – 7.14 (m, 3H), 6.58 (s, 1H), 6.44 (dd, *J* = 16.2, 2.1 Hz, 1H), 6.29 (s, 1H), 4.93 – 4.86 (m, 3H), 3.80 (s, 3H), 3.23 (s, 3H), 3.04 (s, 1H), 2.90 (dd, *J* = 15.5, 3.9 Hz, 1H), 2.72 – 2.63 (m, 3H), 2.37 – 2.30 (m, 3H), 2.06 – 1.96 (m, 3H), 1.91 – 1.30 (m, 11H).

**$^{13}$C NMR (101 MHz, CDCl$_3$)** δ 175.00, 174.61, 168.92, 167.11, 154.18, 141.10, 138.51, 138.05, 135.34, 132.00, 131.77, 129.99, 129.10, 128.62, 128.58, 127.87, 126.27, 125.38, 121.07, 120.14, 111.88, 96.51, 75.97, 52.07, 50.75, 49.85, 36.70, 36.61, 35.75, 35.23, 34.01, 33.71, 32.74, 32.34, 31.70, 31.64, 27.07, 26.28, 25.94.

**MS (ES+):** m/z calc. for $C_{43}H_{48}ClN_3O_9$: 785.3; found: 808.8 $[M+Na]^+$.

<u>Compound **12**</u>

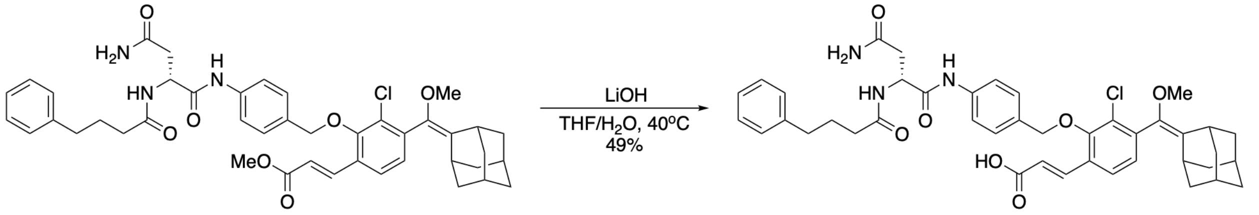

**10** **12**

Compound **10** (23 mg, 0.030 mmol, 1 eq.) was dissolved in 1 mL of 4:1 THF/$H_2O$. LiOH (7.3 mg, 0.305 mmol, 10 eq.) were added and the reaction was heated to 40°C. The reaction was monitored by RP-HPLC (70-100% ACN in water, 0.1% TFA). Upon completion, the solvent was evaporated under reduced pressure. The product was purified by preparative RP-HPLC (70-100% ACN in water, 0.1% TFA) to afford compound **12** as a white solid (11 mg, 49% yield).

**$^{1}$H NMR (400 MHz, MeOD)** δ 7.93 – 7.86 (m, 1H), 7.61 – 7.54 (m, 3H), 7.37 (d, *J* = 8.6 Hz, 2H), 7.27 – 7.21 (m, 2H), 7.19 – 7.12 (m, 3H), 7.09 (d, *J* = 8.0 Hz, 1H), 6.47 (d, *J* = 16.1 Hz, 1H), 5.04 – 4.95 (m, 2H), 3.29 (s, 3H), 3.24 (s, 1H), 2.85 – 2.60 (m, 4H), 2.29 (t, *J* = 7.5 Hz, 2H), 2.03 (s, 1H), 1.97 – 1.75 (m, 14H).

**$^{13}$C NMR (101 MHz, MeOD)** δ 175.99, 174.79, 171.50, 169.97, 154.72, 142.98, 141.01, 140.01, 139.10, 133.10, 132.95, 131.39, 130.98, 130.84, 129.53, 129.38, 129.05, 126.92, 126.41, 121.68, 121.11, 76.79, 57.39, 52.36, 40.11, 40.01, 39.65, 39.57, 38.13, 37.99, 36.19, 34.39, 31.06, 29.83, 29.70, 28.60.
**MS (ES-):** m/z calc. for $C_{42}H_{46}ClN_3O_7$: 739.3; found: 738.8.

<u>Probe **4**</u>

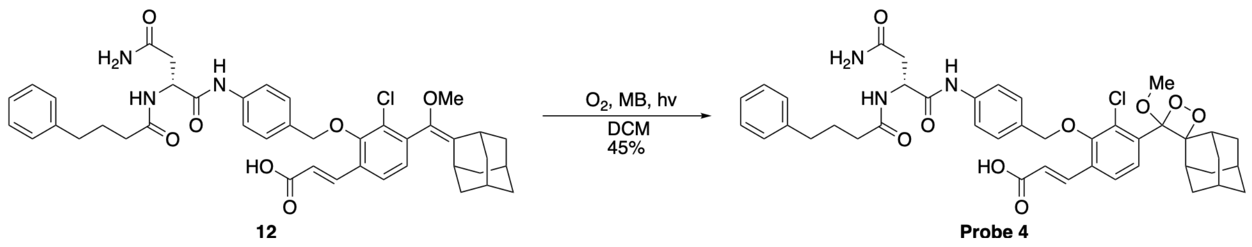

Compound **12** (11 mg, 0.015 mmol) and a catalytic amount of methylene blue were dissolved in 10 mL of DCM. Then, oxygen was bubbled through the solution while irradiating with yellow light. The reaction was monitored by RP-HPLC (70-100% ACN in water, 0.1% TFA). Upon completion, the solvent was evaporated under reduced pressure. The crude product was purified by preparative RP-HPLC (70-100% ACN in water, 0.1% TFA) to afford **Probe 4** as a white solid (5.2 mg, 45% yield).

**$^{1}$H NMR (400 MHz, DMSO)** δ 10.03 (s, 1H), 8.10 (d, *J* = 7.6 Hz, 1H), 7.92 (d, *J* = 8.2 Hz, 1H), 7.80 – 7.70 (m, 2H), 7.64 – 7.56 (m, 2H), 7.35 (d, *J* = 7.7 Hz, 2H), 7.30 (s, 1H), 7.27 – 7.22 (m, 2H), 7.18 – 7.14 (m, 3H), 6.89 (s, 1H), 6.62 (d, *J* = 16.1 Hz, 1H), 4.86 (q, *J* = 10.6 Hz, 2H), 4.68 (dd, *J* = 14.2, 6.7 Hz, 1H), 3.09 (s, 3H), 2.86 (s, 1H), 2.58 – 2.50 (m, 4H), 2.45 – 2.39 (m, 1H), 2.21 (d, *J* = 12.2 Hz, 1H), 2.13 (t, *J* = 7.3 Hz, 2H), 1.89 (s, 1H), 1.81 – 1.51 (m, 9H), 1.44 (d, *J* = 12.3 Hz, 1H), 1.31 (d, *J* = 12.8 Hz, 1H), 1.21 – 1.14 (m, 1H).

**$^{13}$C NMR (101 MHz, DMSO)** δ 172.04, 171.19, 170.22, 167.14, 153.57, 141.85, 139.37, 136.75, 134.05, 131.37, 130.30, 129.37, 128.37, 128.28, 126.10, 125.73, 122.99, 119.09, 111.21, 95.43, 75.49, 50.67, 49.41, 37.13, 35.89, 34.57, 33.29, 33.07, 31.86, 31.67, 31.10, 30.87, 27.08, 25.52, 25.18.

**MS (ES-):** m/z calc. for $C_{42}H_{46}ClN_3O_9$: 771.3; found: 884.9 $[M+TFA-H]^-$.

### Methods

##### Activity measurements with purified ClbP

Probes **1–4** were prepared as 10 mM stock solutions in DMSO, aliquoted and kept frozen (–20 °C). Aliquots were thawed at room temperature, protected from light and any leftover solution was discarded once thawed. To obtain the working solution, 1 µL of the probe stock was added to 1 mL of PBS containing 10% DMSO. The probes were pre-incubated for 30 minutes, after which 99 µL of each solution was transferred to a 96-well plate. Stocks of WT or S95A ClbP, prepared as previously described,[4] were thawed on ice and diluted in PBS to final concentrations ranging from 13 nM to $10^{-8}$ nM. Subsequently, 1 µL of enzyme was added to the assay mixture, and the measurement was initiated in the microplate reader for chemiluminescence measurements (probes **2-4**, DMSO 10% v/v, temperature 27 °C) or fluorescence (probe **1**, DMSO 10% v/v, temperature 27 °C). Total light emission was calculated for 40 minutes.

##### Activity measurements with purified ZmaM

Stocks of ClbP were thawed on ice and diluted in PBS with 5 mM $MgSO_4$ and 1mM ATP to provide a solution containing 62.5 nM of purified enzyme. Separately, a suspension of probe **2** (10 mM in DMSO) was diluted into PBS (to 50 µM) and incubated at room temperature for an hour. To initiate the assay, the solution of probe **2** (10 µL) was added to the enzyme solution (40 µL, with final concentrations of 10 µM probe **2** and 50 nM ZmaM) and incubated in the microplate reader for chemiluminescence measurements (temperature 37 °C, gain 135, read height 4.50 mm, integration time 20 sec, interval 1 minute).

##### Measurements of ClbP inhibitor activity

Stocks of ClbP were thawed on ice and diluted in PBS to provide a solution containing 25 nM of purified enzyme. The ClbP pinacol boronic acid inhibitor (**13**) was prepared as previously described[5] and dissolved in DMSO (10 mM). The stock solution of **13** was diluted in PBS to the desired concentration ranges (20 µL, for a final concentration ranging from 12.5 µM to 0.01 nM) and was incubated with purified ClbP (20 µL, for a final concentration of 10 nM) for 15 minutes. Separately, a suspension of probe **2** (10 mM in DMSO) was diluted into PBS (to 50 µM) and incubated at room temperature for an hour. To initiate the assay, the solution of probe **2** (10 µL, to provide a final concentration of 10 nM) was added to the enzyme / inhibitor solution and incubated in the microplate reader for chemiluminescence measurements (temperature 37 °C, gain 135, read height 4.50 mm, integration time 20 sec, interval 1 minute). Total light emission was calculated after 15 minutes and fitted through a non-linear curve to determine the $IC_{50}$ value.

##### Measurement of ClbP activity with competitive substrate S1

Stocks of WT or S95A ClbP solution were thawed on ice. Molecule **S1** was prepared as previously described[4] and dissolved in DMSO (1 mM). ClbP samples (diluted in PBS) were treated either with DMSO or a solution of molecule **S1**, to a final concentration of 1.25% v/v DMSO and 12.5 nM of enzyme and left to stand at room temperature for 15 minutes. Separately, a suspension of probe **2** (10 mM in DMSO) was diluted into PBS (to 50 µM) and incubated at room temperature for an hour. To initiate an assay, the probe **2** solution (10 µL) was added to the enzyme solution (40 µL, with final concentrations of 10 µM probe **2** and 10 nM ClbP) and placed in the microplate reader for chemiluminescence measurements (temperature 37 °C, gain 135, read height 4.50 mm, integration time 20 sec, interval 1 minute).

##### Measurements of probe activity with *E. coli* BW25113 BAC-*pks* or BAC-*pksΔclbP*

One 5 mL starter culture of *E. coli* BW25113 WT or *E. coli* BW25113 *ΔclbP* was inoculated from a frozen glycerol stock into LB medium supplemented with chloramphenicol (35 µg/mL) and grown aerobically overnight with shaking (37 °C, 200 rpm). The overnight cultures were diluted 1:1000 into M9 medium supplemented with chloramphenicol (35 µg/mL) and grown aerobically overnight with shaking (37 °C, 200 rpm). Cultures were pelleted (10 minutes at 3200 x *g*), resuspended in PBS to an $OD_{600}$ of 0.4. Separately, suspensions of probes **1-4** (10 mM in DMSO) were diluted into PBS (to 50 µM) and left to stand at room temperature for an hour. To initiate an assay, 40 µL of the bacterial culture and 10 µL of the probe solution (to provide a final concentration of 10 µM) were combined into a 384-well plate and placed in the microplate reader for chemiluminescence measurements (probes **2**-**4**, temperature 37 °C, linear shake 1:30 min, gain 135, read height 4.50 mm, integration time 20 sec, interval 5 minutes) or fluorescence (probe **1**, temperature 37 °C, linear shake 1:30 min, gain 35, read height 1.00 mm, excitation filter 360/40 nm, emission filter 440/30 nm, interval 5 minutes). Total light emission was calculated after one hour.

##### RT-qPCR of ClbP transcripts with *E. coli* BW25113 BAC-*pks* or BAC-*pksΔclbP*

One 5 mL starter culture of *E. coli* BW25113 WT or *E. coli* BW25113 *ΔclbP* was inoculated from a frozen glycerol stock into LB medium supplemented with chloramphenicol (35 µg/mL) and grown aerobically overnight with shaking (37 °C, 200 rpm). The overnight cultures were diluted 1:1000 into M9 medium (50 mL each) supplemented with chloramphenicol (35 µg/mL) and grown aerobically overnight with shaking (37 °C, 200 rpm). Cultures were pelleted (10 minutes at 3200 x *g*), resuspended in PBS to an $OD_{600}$ of 0.4. Separately, suspensions of probe **4** (10 mM in DMSO) were diluted into PBS (to 50 μM) and left to stand at room temperature for an hour. To initiate an assay, 24.5 mL of the WT bacterial culture or 4.9 mL of the ΔClbP bacterial culture and 500 or 100 μL of the probe solution, respectively (to provide a final concentration of 1 μM), were combined into 50 mL falcon tubes and placed in an incubator with shaking (37 °C, 200 rpm). A 5 mL sample was collected immediately from each replicate for T = 0 min, pelleted (3 minutes at 3000 x *g*), resuspended in TRIzol reagent (Invitrogen, catalog number 15596-026), and frozen immediately at –80 °C. Samples were collected again at T = 30 min, 60 min and 120 min for the WT bacteria, and at 120 min for the ΔClbP bacteria. Total RNA was extracted with the Zymo Research Direct-Zol RNA MiniPrep plus kit (R2070) according to the manufacturer's recommendations. From the total RNA, 50ng were used per sample for complementary DNA synthesis and PCR amplification using the Luna Universal One-Step RT-qPCR Kit according to the manufacturer's protocol (NEB, E30005S). The following primers were used for *gapA* and *clbP* amplification: gapA-FWD: 5'-TCAGAAAACCGTTGATGGCCC-3'; gapA-REV: 5'-ACGCCATACCAGTCAGTTTGC-3'; clbP-FWD: 5'-AGCCTTTCTGTGCAACCAGG-3'; clbP-REV: 5'-ATTGTCAACAAGGCAAGCGG-3'. Assays were performed using a Bio-Rad CFX Opus Real-Time PCR System, and fold changes in transcript amounts were calculated by the $\Delta\Delta C_T$ method normalized to one replicate at T = 0 min.

#### Measurements of probe LOD with *E. coli* Nissle 1917

One 5 mL starter culture of *E. coli* Nissle 1917 was inoculated from a frozen glycerol stock into LB medium and aerobically grown overnight with shaking (37 °C, 200 rpm). The overnight cultures were diluted 1:1000 into M9 medium and grown aerobically overnight with shaking (37 °C, 200 rpm). Cultures were pelleted (10 minutes at 3200 x *g*), resuspended in PBS to an $OD_{600}$ of 1.0. Actual CFU was determined to be ~ 5 x $10^8$ CFU/mL by plating serial dilutions. Separately, suspensions of probes **1-4** (10 mM in DMSO) were prepared by diluting into PBS (to 500 μM for probe **1**, 50 μM for probes **2**-**4**) and let them stand at room temperature for an hour. Serial dilutions of the *E. coli* Nissle 1917 were prepared in PBS by combining 40 μL of the bacterial culture suspensions with 10 μL of the probe solution into a 384-well plate (to a final concentration of 100 μM for probe **1**, 10 μM for probes **2**-**4**) and place in the microplate reader for chemiluminescence measurements (probes **2**-**4**, temperature 37 °C, linear shake 1:30 min, gain

135, read height 4.50 mm, integration time 20 sec, interval 5 minutes) or fluorescence (probe **1**, temperature 37 °C, linear shake 1:30 min, gain 35, read height 1.00 mm, excitation filter 360/40 nm, emission filter 440/30 nm, interval 5 minutes). Total light emission was calculated after one hour.

#### Measurements of probe activity with native *pks*⁺ *E. coli* isolates

A 5 mL starter culture of *pks*⁺ *E. coli* (CFT073, CCR20, M1/5, ATCC25922 and SP15) was inoculated from a frozen glycerol stock into LB medium and grown aerobically overnight with shaking (37 °C, 200 rpm). The overnight cultures were diluted 1:1000 into M9 medium and grown overnight with shaking (37 °C, 200 rpm). Cultures were pelleted (10 minutes at 3200 x *g*), and resuspended in PBS to an $OD_{600}$ of 0.6. Separately, suspensions of probes **1-4** (10 mM in DMSO) were prepared by diluting into PBS (to 50 µM) and left them to stand at room temperature for an hour. To initiate an assay, 40 µL of the bacterial culture suspensions with 10 µL of the probe solution (to a final probe concentration of 10 µM) were combined into a 384-well plate and placed in the microplate reader for chemiluminescence (probes **2**-**4**, temperature 37 °C, linear shake 1:30 min, gain 135, read height 4.50 mm, integration time 20 sec, interval 5 minutes) or fluorescence (probe **1**, temperature 37 °C, linear shake 1:30 min, gain 35, read height 1.00 mm, excitation filter 360/40 nm, emission filter 440/30 nm, interval 5 minutes). Total light emission was calculated after one hour.

#### Measurements of ClbP inhibition in cultures of *E. coli* Nissle 1917

A 5 mL starter culture of *E. coli* Nissle 1917 was inoculated from a frozen glycerol stock into LB medium and grown aerobically overnight with shaking (37 °C, 200 rpm). The overnight culture was diluted 1:1000 into M9 medium and grown aerobically overnight with shaking (37 °C, 200 rpm). Cultures were pelleted (10 minutes at 3200 x *g*), resuspended in PBS to an $OD_{600}$ of 0.1. Resuspended cells (39 µL) were treated either with DMSO (1 µL) or a solution of Inhibitor **13** (1 mM in DMSO, 1 µL) and incubated for an hour at room temperature. Separately, suspensions of probe **2**-**4** (10 mM in DMSO) were prepared by diluting into PBS (to 50 µM) and left them to stand at room temperature for an hour. To initiate an assay, the bacterial suspension (40 µL) was combined with 10 µL of the probe solution (to a final concentration of inhibitor **13** of 20 µM, and a probe concentration of 10 µM) into a 384-well plate and placed in the microplate reader for chemiluminescence measurements (probe **2**-**4**, temperature 37 °C, linear shake 1:30 min, gain 135, read height 4.50 mm, integration time 20 sec, interval 5 minutes). Total light emission was calculated after one hour.

#### Measurements of probe activity with other Gram-negative bacteria

A 5 mL starter culture of *Klebsiella pneumoniae* strains WGLW3 and WGLW5, *Erwinia oleae* DAPP-PG-531 and *Pseudovibrio denitrificans* JCM 12308 was inoculated from a frozen glycerol stock into NB medium (for *K. pneumoniae*), TSB medium (for *E. oleae*) or MB medium (for *P. denitrificans*) and grown aerobically overnight with shaking (37 °C, 200 rpm for *K. pneumoniae,* and 28 °C, 200 rpm for *E. oleae* and *P. denitrificans*). The overnight cultures were diluted 1:1000 into M9 medium (for *K. pneumoniae*), TSB medium (for *E. oleae*) or MB medium (for *P. denitrificans*) and grown overnight with shaking (37 °C, 200 rpm for *K. pneumoniae,* and 28 °C, 200 rpm for *E. oleae* and *P. denitrificans*). Cultures were pelleted (10 minutes at 3200 x *g*), and resuspended in PBS to an $OD_{600}$ of 0.5. Separately, suspensions of probes **2-4** (10 mM in DMSO) were prepared by diluting into PBS (to 50 µM) and left them to stand at room temperature for an hour. To initiate an assay, 40 µL of the bacterial culture suspensions with 10 µL of the probe solution (to a final probe concentration of 10 µM) were combined into a 384-well plate and placed in the microplate reader for chemiluminescence (probes **2**-**4**, temperature 37 °C, linear shake 1:30 min, gain 135, read height 4.50 mm, integration time 20 sec, interval 5 minutes).

#### Measurements of ClbP inhibition in other Gram-negative bacteria

A 5 mL starter culture of *Klebsiella pneumoniae* strain WGLW3, *Erwinia oleae* DAPP-PG-531 and *Pseudovibrio denitrificans* JCM 12308 was inoculated from a frozen glycerol stock into NB medium (for *K. pneumoniae*), TSB medium (for *E. oleae*) or MB medium (for *P. denitrificans*) and grown aerobically overnight with shaking (37 °C, 200 rpm for *K. pneumoniae,* and 28 °C, 200 rpm for *E. oleae* and *P. denitrificans*). The overnight cultures were diluted 1:1000 into M9 medium (for *K. pneumoniae*), TSB medium (for *E. oleae*) or MB medium (for *P. denitrificans*) and grown overnight with shaking (37 °C, 200 rpm for *K. pneumoniae,* and 28 °C, 200 rpm for *E. oleae* and *P. denitrificans*). Cultures were pelleted (10 minutes at 3200 x *g*), resuspended in PBS to an $OD_{600}$ of 0.5. Resuspended cells (39 µL) were treated either with DMSO (1 µL) or a solution of Inhibitor **13** (1 mM in DMSO, 1 µL) and incubated for an hour at room temperature. Separately, suspensions of probe **2**-**4** (10 mM in DMSO) were prepared by diluting into PBS (to 50 μM) and left them to stand at room temperature for an hour. To initiate an assay, the bacterial suspension (40 μL) was combined with 10 μL of the probe solution (to a final concentration of inhibitor **13** of 20 μM, and a probe concentration of 10 μM) into a 384-well plate and placed in the microplate reader for chemiluminescence measurements (probe **2**-**4**, temperature 37 °C, linear shake 1:30 min, gain 135, read height 4.50 mm, integration time 20 sec, interval 5 minutes). Total light emission was calculated after one hour.

##### Measurements of probe activity in germ-free stool samples

Mouse stool samples from germ-free C57BL/6 mice that were housed at the Harvard T.H. Chan Gnotobiotics Center for Mechanistic Microbiome Studies in semi-rigid isolators (Plastic Concepts Inc) were provided by Prof. Wendy Garrett. A germ-free stool supernatant was prepared by weighing mouse stool pellets (wet weight), resuspending by pipetting and vortexing to provide a 20 mg/mL slurry in PBS, pelleting debris (1 minute at 300 x *g*), and removing the stool supernatant. For probe stability experiments, the stool supernatant (40 μL) was diluted with PBS into a 384-well microplate to the desired final concentrations (5, 2 or 1 mg/mL) in a 50 μL volume with the optional addition of purified ClbP WT or S95A (to a final concentration of 10 nM). For filtration experiments, the supernatant was diluted to 10 mg/mL and passed through a sterile filter (100 μm, 70 μm or 40 μm, Greiner Bio-One) with the aid of a P1000 pipette. The filtered or unfiltered stool supernatants were dispensed and diluted (to 6.25 mg/mL, for a total volume 40 μL) into a 384-well microplate with the addition of purified ClbP WT (at a final concentration of 10 nM). Separately, a suspension of probe **2** (10 mM in DMSO) was prepared by diluting into PBS (to 50 μM) and left to stand at room temperature for an hour. To initiate an assay, probe **2** solution was added to the stool suspension (to a final probe concentration of 10 μM) and placed in the microplate reader for chemiluminescence measurements (temperature 37 °C, gain 135, read height 4.50 mm, integration time 20 sec, interval 1 minute). Total light emission was calculated after one hour.

##### Measurements of probe activity in stool samples containing *E. coli* BW25113

A 5 mL starter culture of *E. coli* BW25113 WT or *E. coli* BW25113 *ΔclbP* was inoculated from a frozen glycerol stock into LB medium supplemented with chloramphenicol (35 μg/mL) and grown aerobically overnight with shaking (37 °C, 200 rpm). The overnight cultures were diluted 1:1000 into M9 medium supplemented with chloramphenicol (35 μg/mL) and grown aerobically overnight with shaking (37 °C, 200 rpm). Cultures were pelleted (10 minutes at 3200 x *g*) and resuspended in PBS to an $OD_{600}$ of 1.0. Separately, suspensions of probes **1-4** (10 mM in DMSO) were diluted into PBS (to 50 μM) and left to stand at room temperature for an hour. The germ-free stool suspension was prepared by weighing wet mouse stool pellets, resuspending to a 10 mg/mL slurry in PBS by pipetting and vortexing, pelleting debris (1 minute at 300 x *g*), and removing the stool supernatant. To initiate an assay, 15 μL of the bacterial suspension, 25 μL of stool solution, and 10 μL of the probe solution were combined (to a final probe concentration of 10 μM) into a 384-well plate and placed in the microplate reader for chemiluminescence measurements (probes **2**-**4**, temperature 37 °C, linear shake 1:30 min, gain 135, read height 4.50 mm, integration time 20 sec, interval 5 minutes) or fluorescence (probe **1**, temperature 37 °C, linear shake 1:30 min, gain 35, read height 1.00 mm, excitation filter 360/40 nm, emission filter 440/30 nm, interval 5 minutes). Total light emission was calculated after one hour.

##### Measurements of probe activity in stool samples containing native *pks*$^+$ *E. coli* isolates

A 5 mL starter culture of naturally-encoding *pks E. coli* (M1/5, ATCC25922, Nissle 1917 and Nissle 1917 *ΔclbP*) was inoculated from a frozen glycerol stock into anaerobic LB medium and grown anaerobically overnight statically (37 °C). The overnight cultures were diluted 1:1000 into anaerobic NCE medium (supplemented with 3 g/L glycerol, as previously described[6]) and grown overnight statically (37 °C). Cultures were pelleted (10 minutes at 3200 x *g*), resuspended in PBS to an $OD_{600}$ of 0.2. Cultures were calculated to have approximately 1.6 x $10^8$ cells/mL (http://www.agilent.com/store/biocalculators/calcODBacterial.jsp). Separately, anaerobic suspensions of probes **1-4** (10 mM in DMSO) were prepared by diluting into PBS (to 500 μM for probe **1**, 125 μM for probes **2**-**4**) and left to stand at room temperature for an hour. The germ-free and ASF stool suspensions were prepared by weighing wet mouse stool pellets, which were then resuspended to a 10 mg/mL slurry in anaerobic PBS by pipetting and vortexing, the debris was pelleted (1 minute at 300 x *g*) and the supernatant removed. To initiate an assay, 15 μL of the bacterial solution, 25 μL of stool solution and 10 μL of the probe solution were combined (to a final probe concentration of 25μM for probes **2**-**4** and 100 μM for probe **1**) into a 384-well plate. The plate was removed from the anaerobic chamber and placed in the microplate reader for chemiluminescence measurements (probes **2**-**4**, temperature 37 °C, linear shake 1:30 min, gain 135, read height 4.50 mm, integration time 20 sec, interval 5 minutes) or fluorescence (probe **1**, temperature 37 °C, linear shake 1:30 min, gain 35, read height 1.00 mm, excitation filter 360/40 nm, emission filter 440/30 nm, interval 5 minutes). Total light emission was calculated after one hour.

#### Detection of hydrolysis products from chemiluminescent probes

Stocks of WT or S95A ClbP solutions were thawed on ice and diluted in PBS to provide solutions containing 20 nM of purified enzyme. Separately, suspensions of probes **2-4** (10 mM in DMSO) were prepared by diluting into PBS (to 2 mM) and left to stand at room temperature for an hour. To initiate a reaction, the enzyme solution (10 µL), or PBS for the no enzyme control, were combined with the probe solution (to provide final concentrations of 10 nM enzyme and 1 mM probe) and incubated for one hour at room temperature. Each assay mixture was quenched with acetonitrile (180 µL) and stored at –80 °C overnight. Once thawed, the samples were spun (20,000 *g* x 20 min), and the supernatants (50 µL) were transferred into separate tubes (with 450 µL of water). Each solution was filtered through a 0.2 µM nylon filter (Cytiva) and placed into an autosample vial for analysis by LC–MS, using an Agilent Q-TOF 6530 equipped with a Dual AJS ESI source and a Kinetex C8 column (2.6 µm, 100 Å, 100 × 4.6 mm) flowing at a rate of 0.5 mL/min in a column compartment heated to 35 °C. Solution A was $H_2O$ + 0.1% formic acid, and Solution B was MeCN + 0.1% formic acid. The LC method was as follows: 5% Solution B for 5 min, 5–95% Solution B over 15 min, 95% Solution B for 5 min, 95–5% Solution B over 2 min, hold at 5% Solution B for 9 min. The following parameters were used for the Q-TOF: gas temp 275 °C, drying gas 11 L/min, nebulizer 35 psi, sheath gas temp 275 °C, sheath gas flow 11 L/min, VCap 3500 V, and nozzle voltage 500 V. An authentic standard of *N*-myristoyl-D-asparagine was prepared as previously described[7].

### Supplementary Table

| Strain | Genotype | Reference |
|---|---|---|
| ***E. coli BW251113 BAC-pks*** | pBeloBAC11 harboring the complete *pks* island of *E. coli* IHE3034 (under native promoter), $Cm^R$ | Nougayrède, J.P. et al.[8] |
| ***E. coli BW251113 BAC-pksΔclbP*** | pBeloBAC11 harboring a *clbP* deletion in the *pks* island of *E. coli* IHE3034 (under native promoter), $Cm^R$ | Dubois, D. et al.[9] |
| ***E. coli* Nissle 1917** | Native *pks* strain, common probiotic | Scaldaferri, P. et al.[10] |
| ***E. coli* Nissle 1917 ΔclbP** | Red/ET recombination of *clbP,* $Cm^R$ | Bian, X. et al.[11] |
| ***E. coli* CCR20** | Native *pks* strain, from a human colon tumor | Buc, E. et al.[12] |
| ***E. coli* CFT073** | Native *pks* strain, from acute pyelonophritis | Hull, R. et al.[13] |
| ***E. coli* SP15** | Native *pks* strain, from newborn meningitis | Johnson, J et al.[14] |
| ***E. coli* M1/5** | Native *pks* strain, from human stool. | Wallenstein, A. et al.[15] |
| ***E. coli* ATCC 25922** | Native *pks* strain, clinical strain. | ATCC. |
| ***K. pneumoniae* WGLW3** | Native *pks* strain, from human stool. | BEI resources. |
| ***K. pneumoniae* WGLW5** | From mouse stool. | BEI resources. |
| ***E. oleae* DAPP-PG-531** | Native *pks* strain, from olive knots. | Moretti, C. et al.[16] |
| ***P. denitrificans* JCM 12308** | Native *pks* strain, from seawater. | Shieh, W.J. et al.[17] |

**Table S1.** List of bacterial strains used strains in this study.

### Supplementary Figures

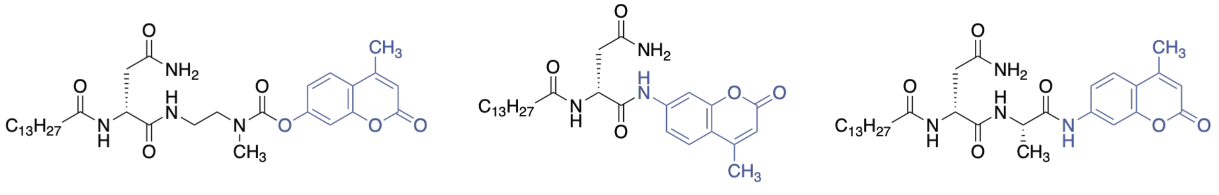

(Volpe et al.) (Hirayama et al.) (Hirayama et al.)

**Figure S1.** Other activity-based fluorescent probes developed for ClbP detection.

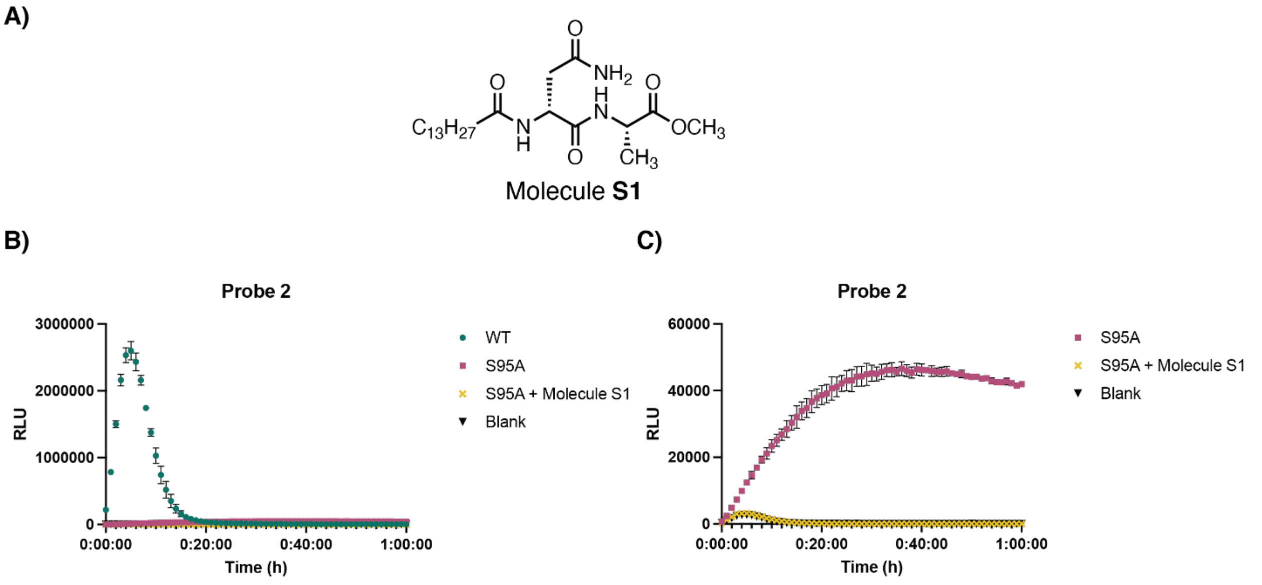

**Figure S2.** Incubation of Probe **2** with a competitive ClbP substrate (**S1**) reduces noise levels for catalytically inactive enzyme. **A)** Structure of the ClbP substrate molecule **S1**. **B)** Luminescent assay with Probe **2** (10 μM), ClbP WT or S95A (10 nM), and optionally molecule **S1** (10 μM) in PBS (37 °C). Error bars represent the ±SD of three biological replicates. **C)** Inset of B), highlighting the ClbP S95A, S95A + **S1** and blank traces.

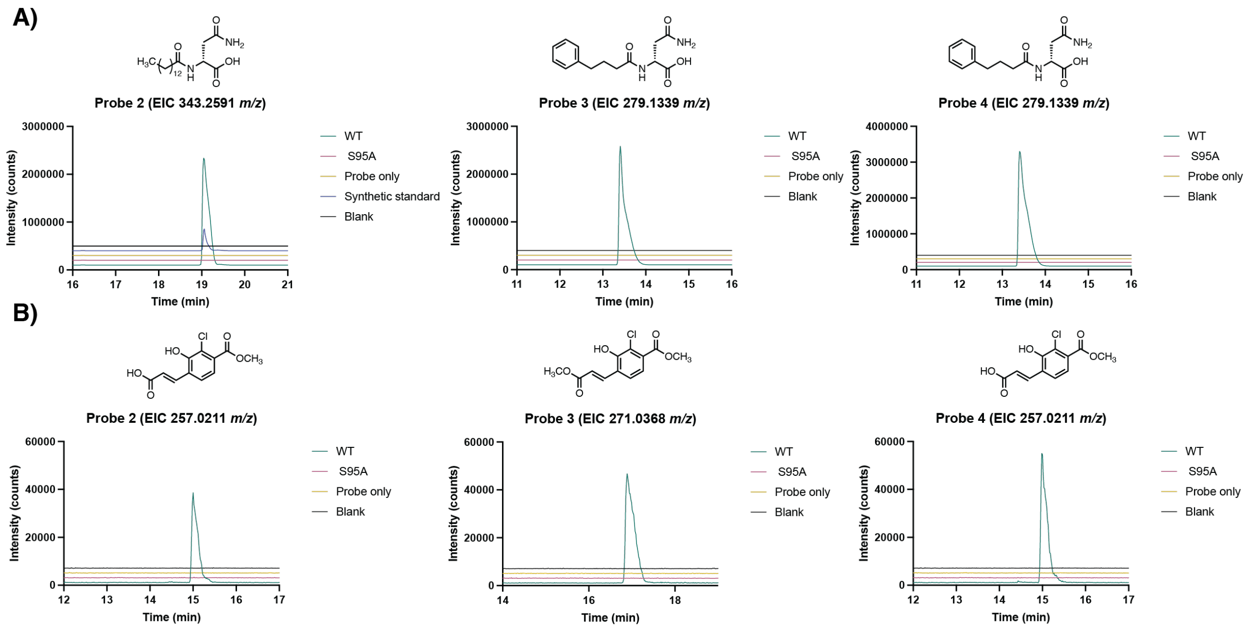

**Figure S3.** LC–MS traces of products obtained after incubation of probes **2-4** with ClbP. **A)** Representative extracted ion chromatograms [M+H$^+$] of prodrug motifs released from probes (three replicates of each). **B)** Representative extracted ion chromatograms [M+H$^+$] of luminophores released from probes (three replicates of each).

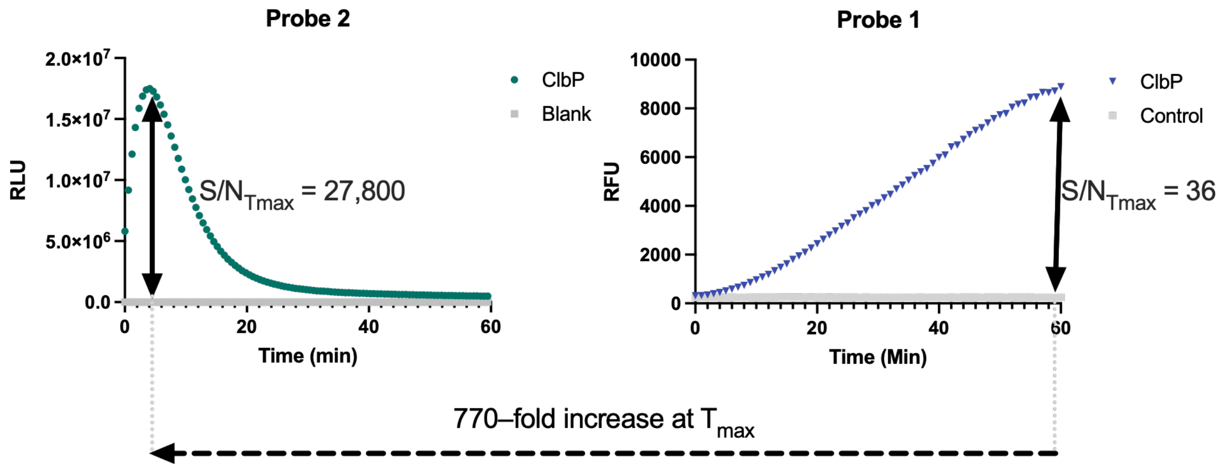

**Figure S4.** Maximum signal-to-noise ratios (S/N$_{Tmax}$) **for chemiluminescent probe 2 (10 µM) and fluorescent probe 1 (10 µM)** with recombinant **ClbP** (13 nM) after incubation for one hour at room temperature in PBS.

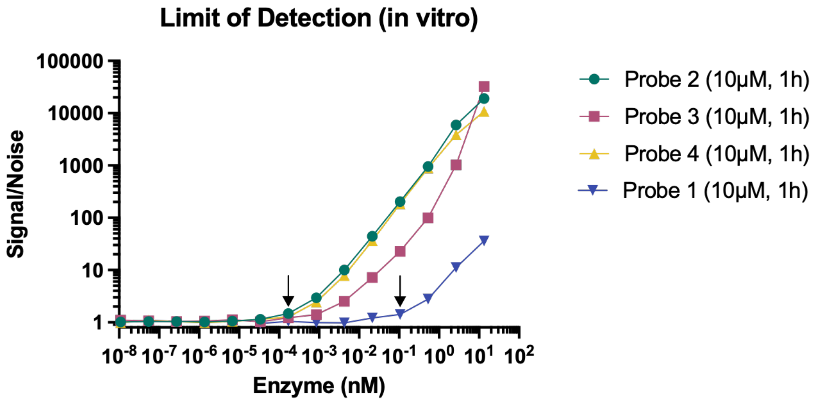

**Figure S5.** Limits of detection **for chemiluminescent probes 2–4 (10 µM) and fluorescent probe 1 (10 µM) with recombinant ClbP** (from $10^{-8}$ nM to 13 nM) after incubation for one hour at room temperature in PBS. Limit of detection are depicted with black arrows in the plot and determined to be at $1.7 \times 10^{-4}$ nM for probes **2-4** and $1.0 \times 10^{-1}$ nM for probe **1**.

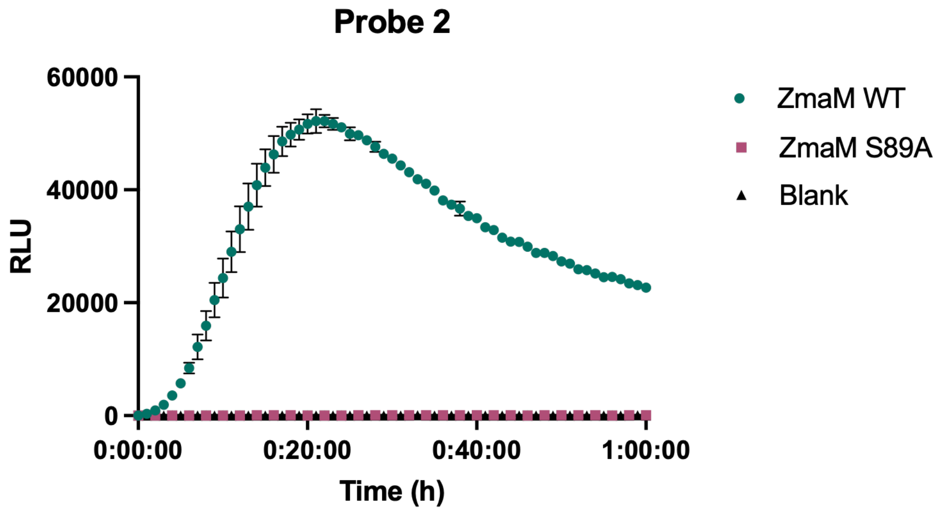

**Figure S6.** Recombinant ZmaM (a ClbP homologue involved in zwittermicin biosynthesis, 27% ID, 50 nM) but not its active site mutant (ZmaM S89A, 50 nM) process chemiluminescent probe **2** (10 µM, 37 °C). Error bars represent the ±SD of three independent measurements.

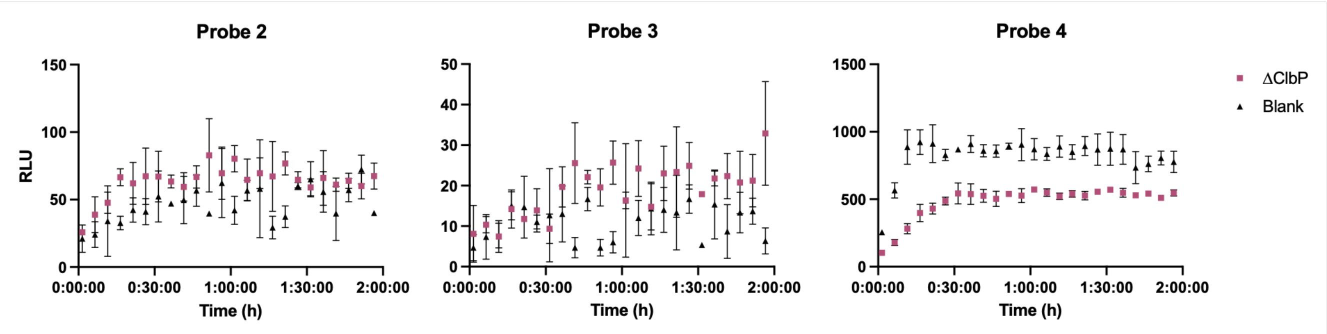

**Figure S7.** Background luminescence levels of *E. coli* BW25113 *ΔclbP* in PBS and blank solutions with chemiluminescent probes **2-4** (10 µM, 37 °C). Error bars represent the ±SD of three biological replicates.

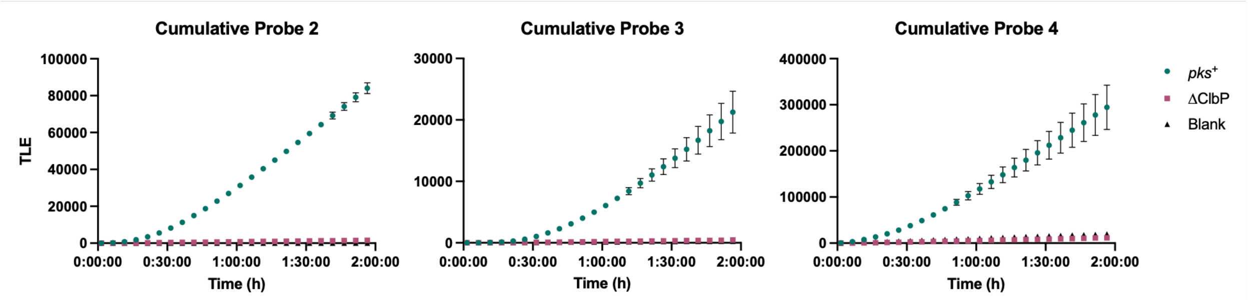

**Figure S8.** Cumulative luminescence levels of *E. coli* BW25113 heterologously expressing the *pks* gene cluster or a *ΔclbP* knockout in PBS with chemiluminescent probes **2-4** (10 μM, 37 °C). Error bars represent the ±SD of three biological replicates.

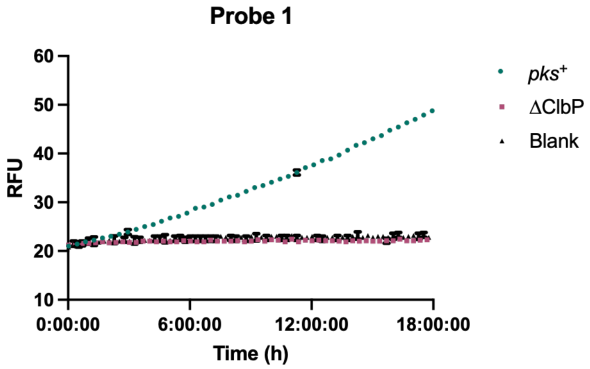

**Figure S9.** Fluorescence levels of *E. coli* BW25113 heterologously expressing the *pks* gene cluster or a *ΔclbP* knockout in PBS with fluorescent probe **1** (10 μM, 37 °C). Error bars represent the ±SD of three biological replicates.

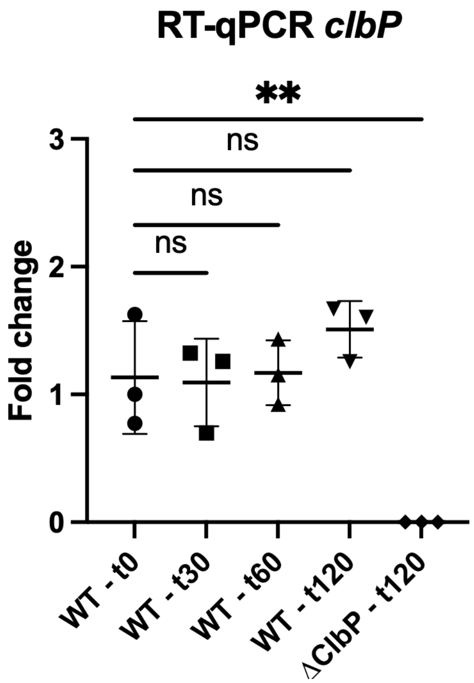

**Figure S10**. RT-qPCR to test the expression level changes of *clbP* during the chemiluminescent assay 0, 30, 60 and 120 minutes in *E. coli* BW 25113 BAC-*pks* and 120 minutes for *E. coli* BW 25113 BAC-*pksΔclbP*. Error bars represent the ±SD of three biological replicates. ***P*<0.01; Not significant *P*>0.05, using one-way ANOVA and Dunnett's multiple comparison test.

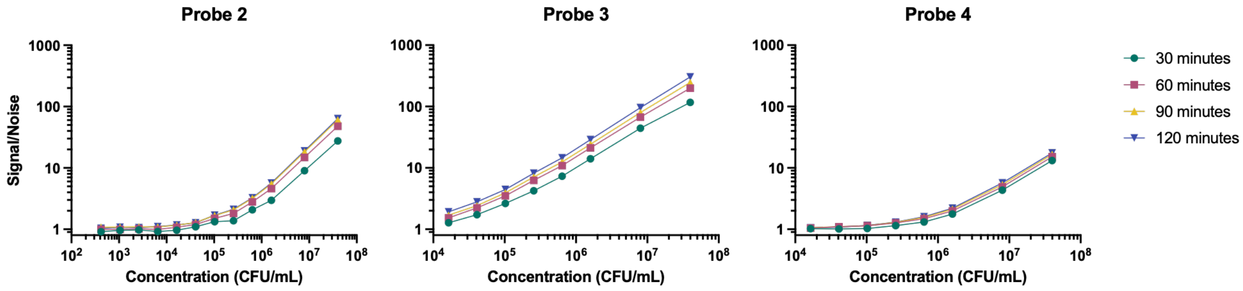

**Figure S11.** Signal-to-noise ratios over different total integration times in the presence of varying amounts of *E. coli* Nissle 1917 for chemiluminescent probes **2**-**4** (10 μM, PBS, 37 °C).

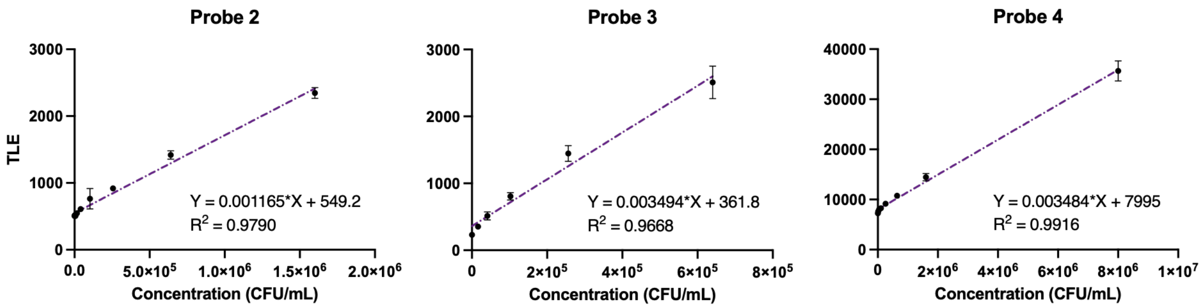

**Figure S12.** Simple linear regression of the amount of *E. coli* Nissle 1917 and total luminescence in the presence of chemiluminescent probes **2**-**4** (10 µM, PBS, 37 °C). Error bars represent the ±SD of three independent measurements.

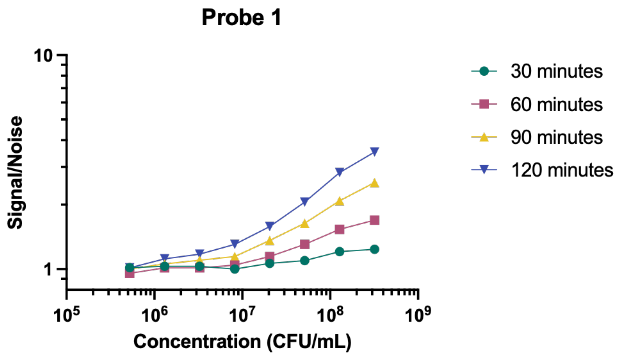

**Figure S13.** Signal-to-noise ratios over different relative times in the presence of varying amounts of *E. coli* Nissle 1917 for fluorescent probe **1** (100 µM, PBS, 37 °C).

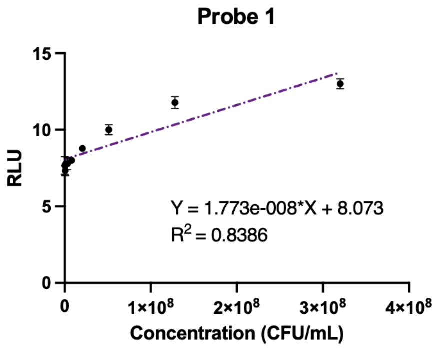

**Figure S14.** Simple linear regression of the amount of *E. coli* Nissle 1917 and relative fluorescence in the presence of fluorescent probe **1** (100 µM, PBS, 37 °C) Error bars represent the ±SD of three biological replicates.

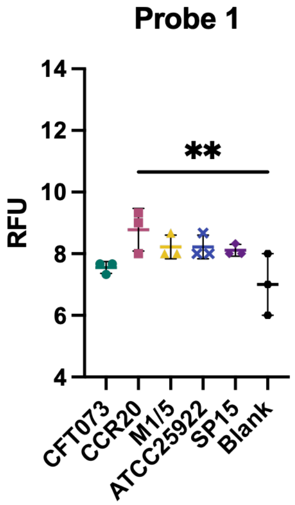

**Figure S15.** Relative fluorescence after one hour incubation of five *pks*⁺ *E. coli* isolates with probe **1** (10 µM, PBS, 37 °C). Error bars represent the ±SD of three biological replicates. Only the shown comparison was found to be significant. ***P*<0.01; Not significant *P*>0.05 using one-way ANOVA and Dunnett's multiple comparison test.

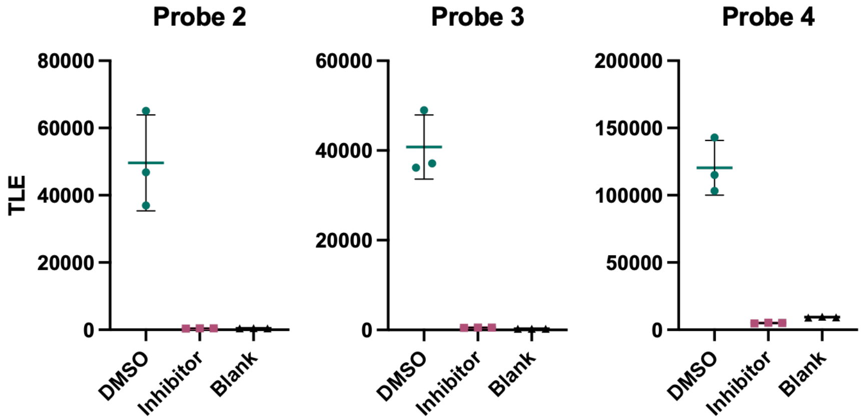

**Figure S16.** Incubation of *E. coli* Nissle 1917 with DMSO (2% v/v) or Inhibitor **13** (20 µM, PBS, 37 °C) with probes **2-4** (10 µM) and light integration after one hour. Error bars represent the ±SD of three biological replicates.

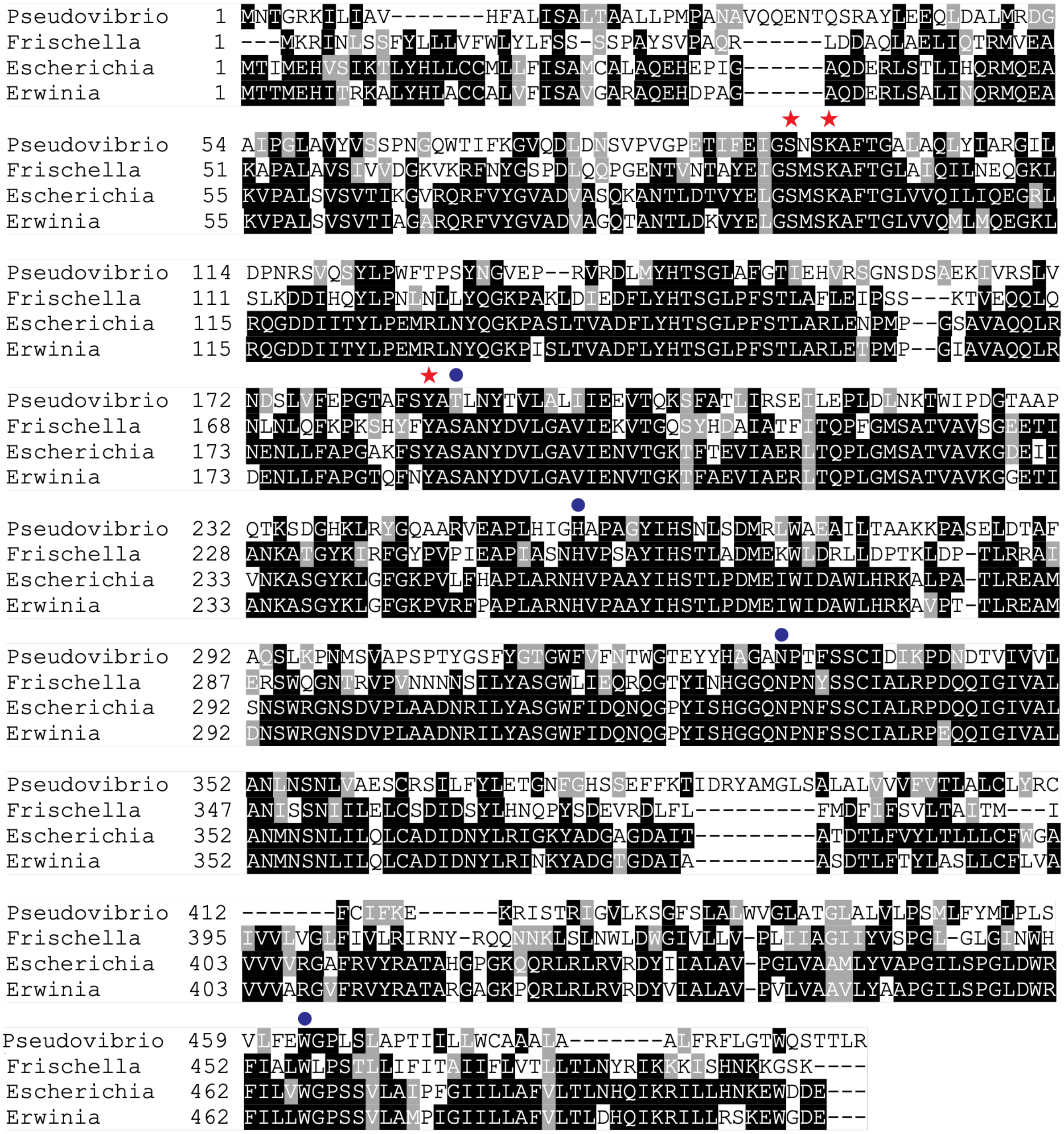

**Figure S17. Multiple sequence alignment of ClbP orthologs.** *Erwinia oleae* (87% ID to *E. coli* ClbP) and *Frischella perrara* (52% ID) have been characterized to produce active genotoxin, whilst *Pseudovibrio denitrificans* (29% ID) bears a colibactin cluster that has not been functionally corroborated. Red stars denote residues involved in catalytic triad, and blue circles represent residues that interact with the prodrug motif in the binding pocket. Sequences were aligned with clustalo[18] (version 1.2.4) with default parameters. MSA figure was created with https://junli.netlify.app/apps/boxshade/.

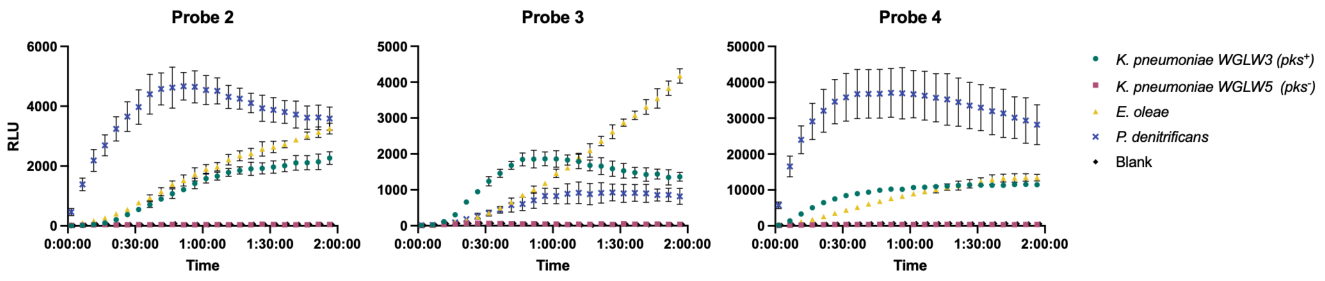

**Figure S18. Chemiluminescent probes are activated by ClbP-encoding Gram-negative bacteria other than *E. coli*.** *Klebsiella pneumoniae* WGLW3 (*pks*+, 100% ID*)*, *K. pneumoniae* WGLW5 (*pks*−), *Erwinia oleae* DAPP-PG-531 (87% ID) and *Pseudovibrio denitrificans* JCM12308 (29 % ID) can process chemiluminescent probes **2**-**4** (10 *µ*M, 37°C, PBS). Error bars represent the ±SD of three biological replicates.

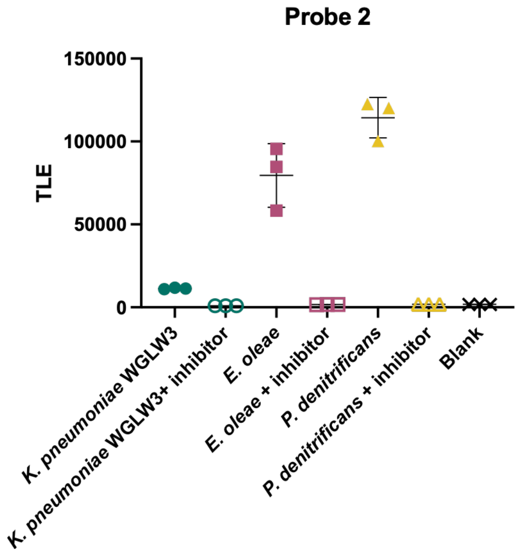

**Figure S19.** Incubation of *K. pneumoniae* WGLW3, *E. oleae* DAPP-PG-531 and *P. denitrificans* JCM13208 with DMSO (2 % v/v) or Inhibitor **13** (20 µM, PBS, 37 °C) with probe **2** (10 µM) and light integration after one hour. Error bars represent the ±SD of three biological replicates.

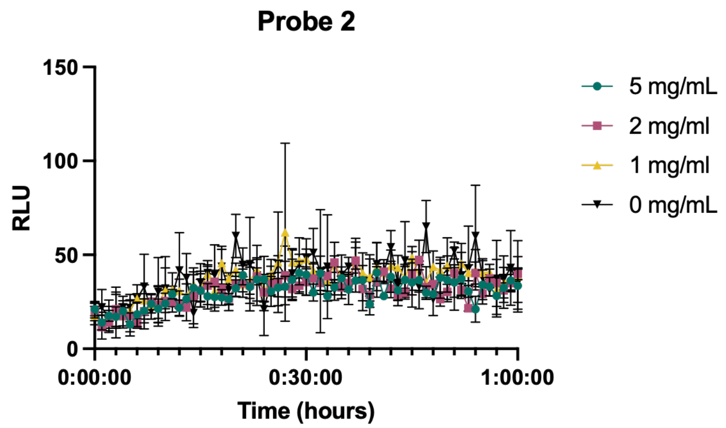

**Figure S20.** Luminescence observed upon incubation of probe **2** (10 µM, PBS, 37 °C) with varying amounts of germ-free stool resuspended in PBS. Error bars represent the ±SD of three biological replicates.

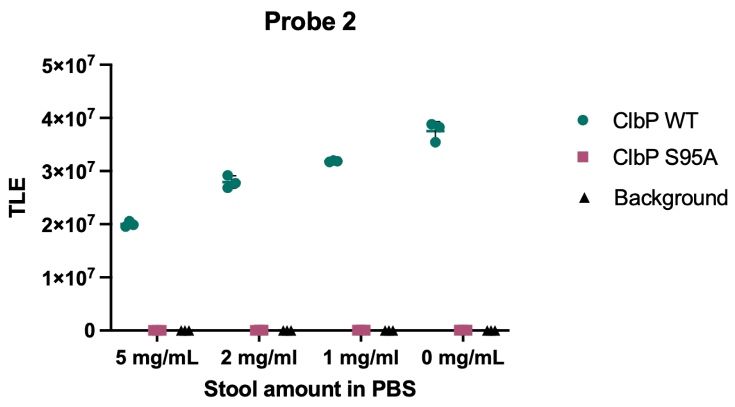

**Figure S21**. Total luminescence observed upon incubation of probe **2** (10 µM, 37 °C) with purified ClbP WT or S95A (10 nM) and varying amounts of germ-free stool resuspended in PBS and light integration after one hour. Error bars represent the ±SD of three biological replicates.

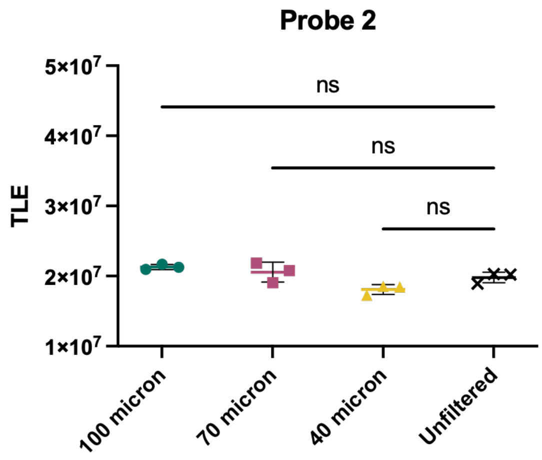

**Figure S22.** **Total luminescence observed upon incubation of probe 2 (10 µM, 37 °C) with purified ClbP WT (10 nM) and filtered or unfiltered germ-free stool resuspensions in PBS (5 mg/mL) and light integration after one hour.** Error bars represent the ±SD of three biological replicates. Not significant *P*>0.05 using one-way ANOVA and Dunnett's multiple comparison test.

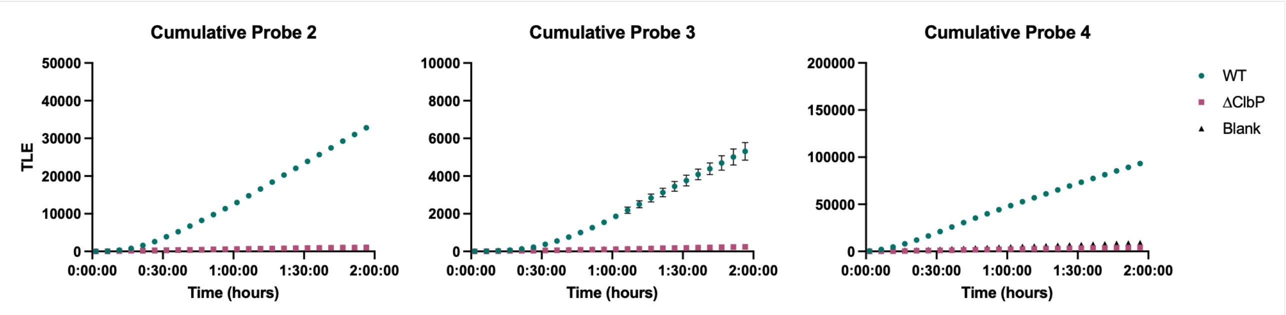

**Figure S23. Cumulative luminescence levels of *E. coli* BW25113 heterologously expressing the *pks* gene cluster or a *ΔclbP* knockout in a germ-free stool resuspension in PBS (5 mg/mL) with chemiluminescent probes 2-4** (10 µM, PBS, 37 °C). Error bars represent the ±SD of three biological replicates.

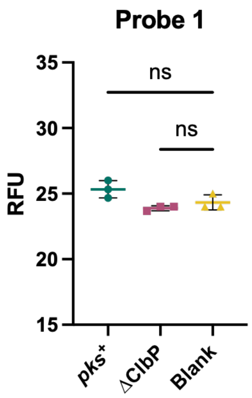

**Figure S24. Relative fluorescence levels of *E. coli* BW25113 heterologously expressing the *pks* gene cluster or a Δ*clbP* knockout in a germ-free stool resuspension in PBS (5 mg/mL) after one hour (10 µM, PBS, 37 ºC).** Error bars represent the ±SD of three biological replicates. Not significant (ns) P>0.05 using one-way ANOVA and Dunnett's multiple comparison test.

### Spectral Data of Probes 2 – 4

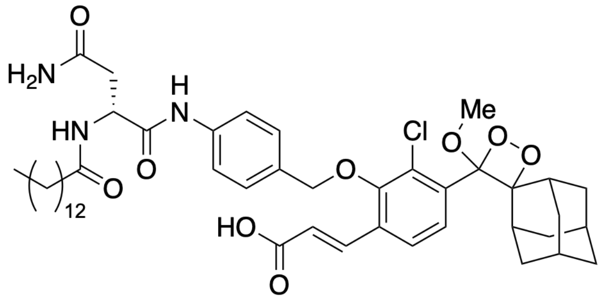

**Probe 2**

Chemical Formula: $C_{46}H_{62}ClN_3O_9$
Exact Mass: 835.42

[1]H NMR of Probe **2**

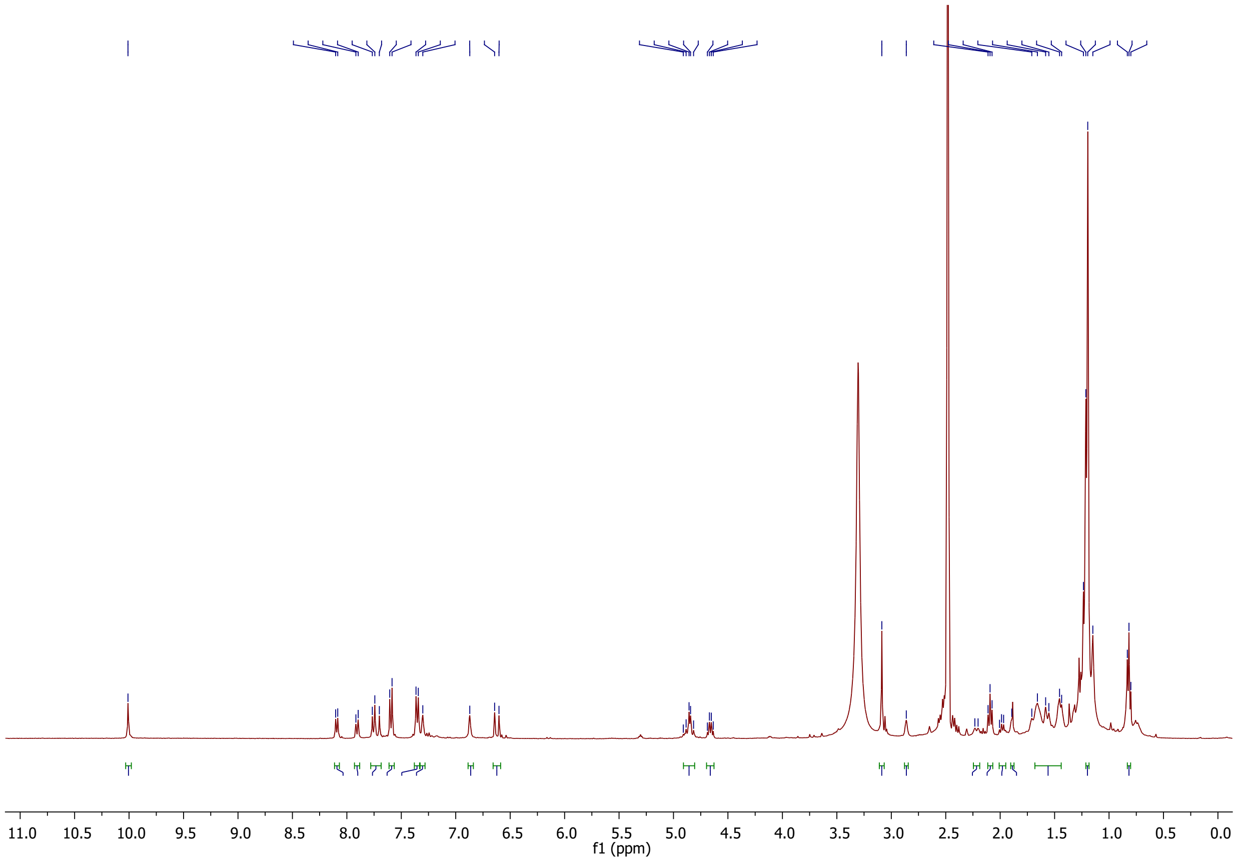

#### $^{13}C$ NMR of Probe **2**

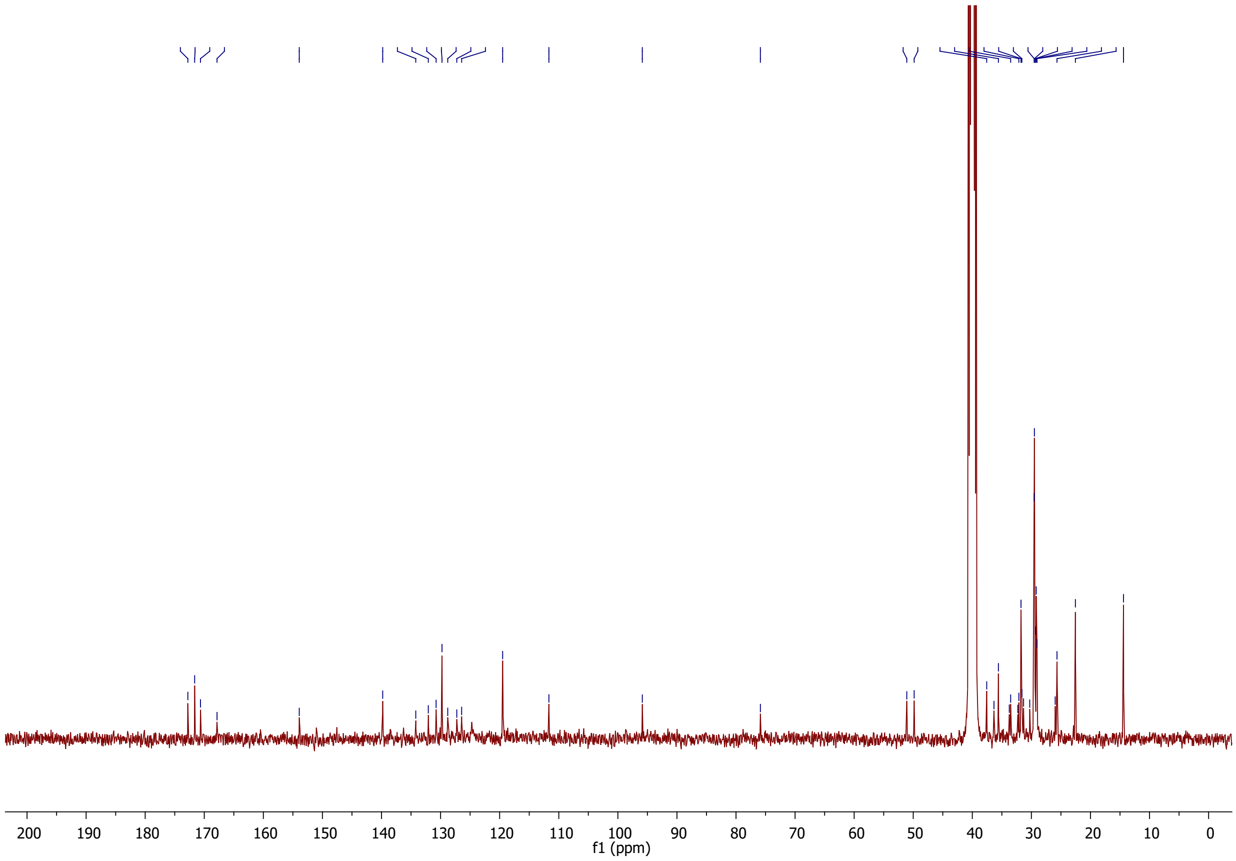

#### MS of Probe **2** (ESI Negative)

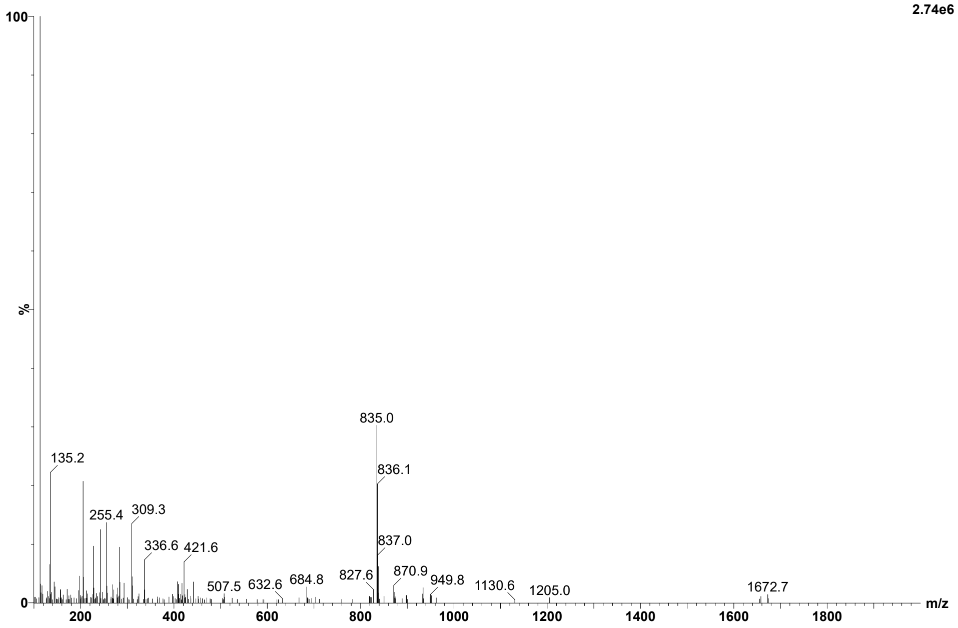

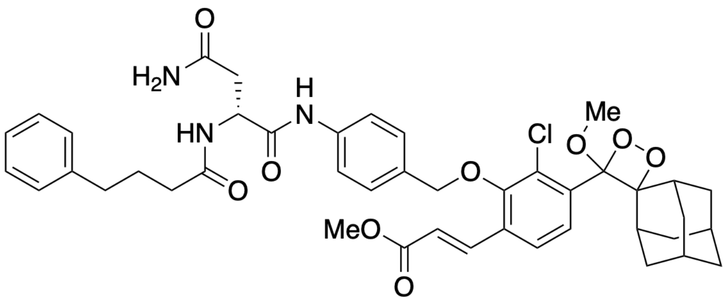

**Probe 3**

Chemical Formula: $C_{43}H_{48}ClN_3O_9$
Exact Mass: 785.31

[1]H NMR of Probe **3**

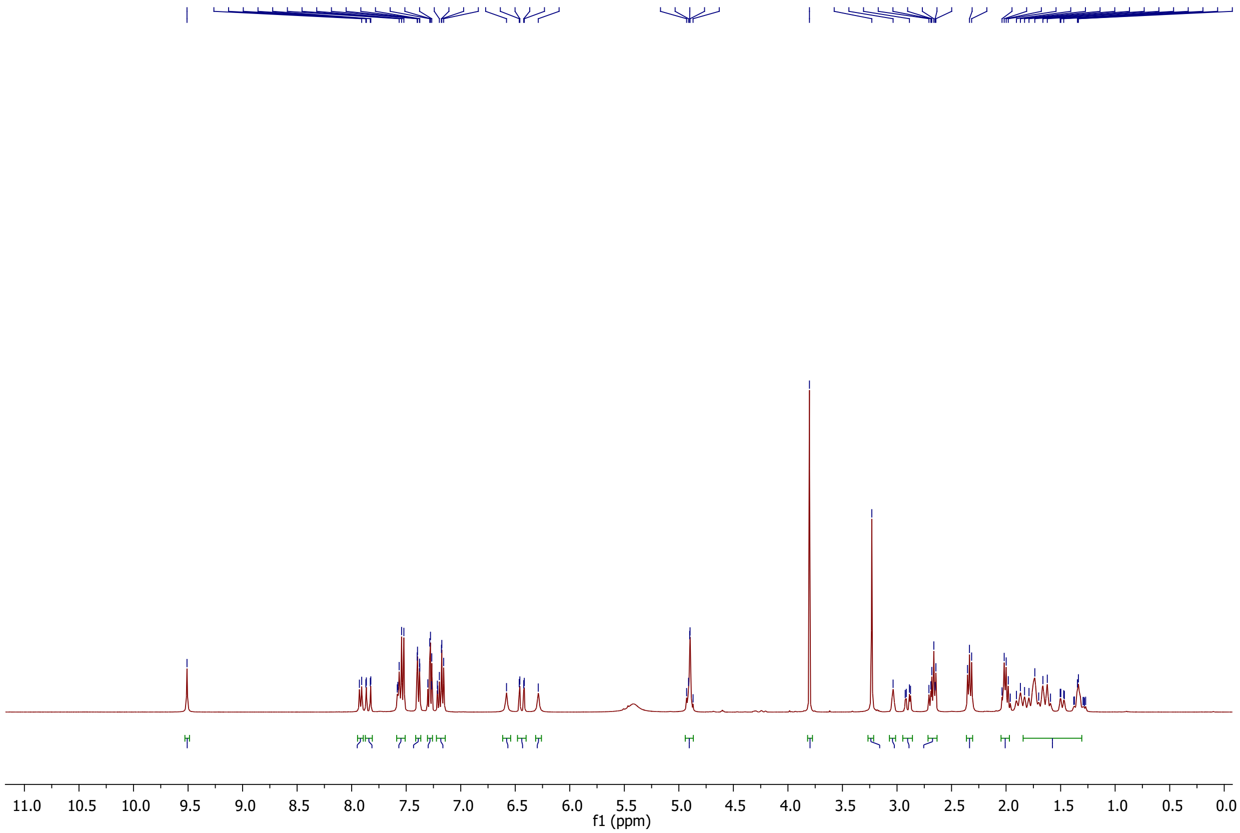

<u>$^{13}$C NMR of Probe **3**</u>

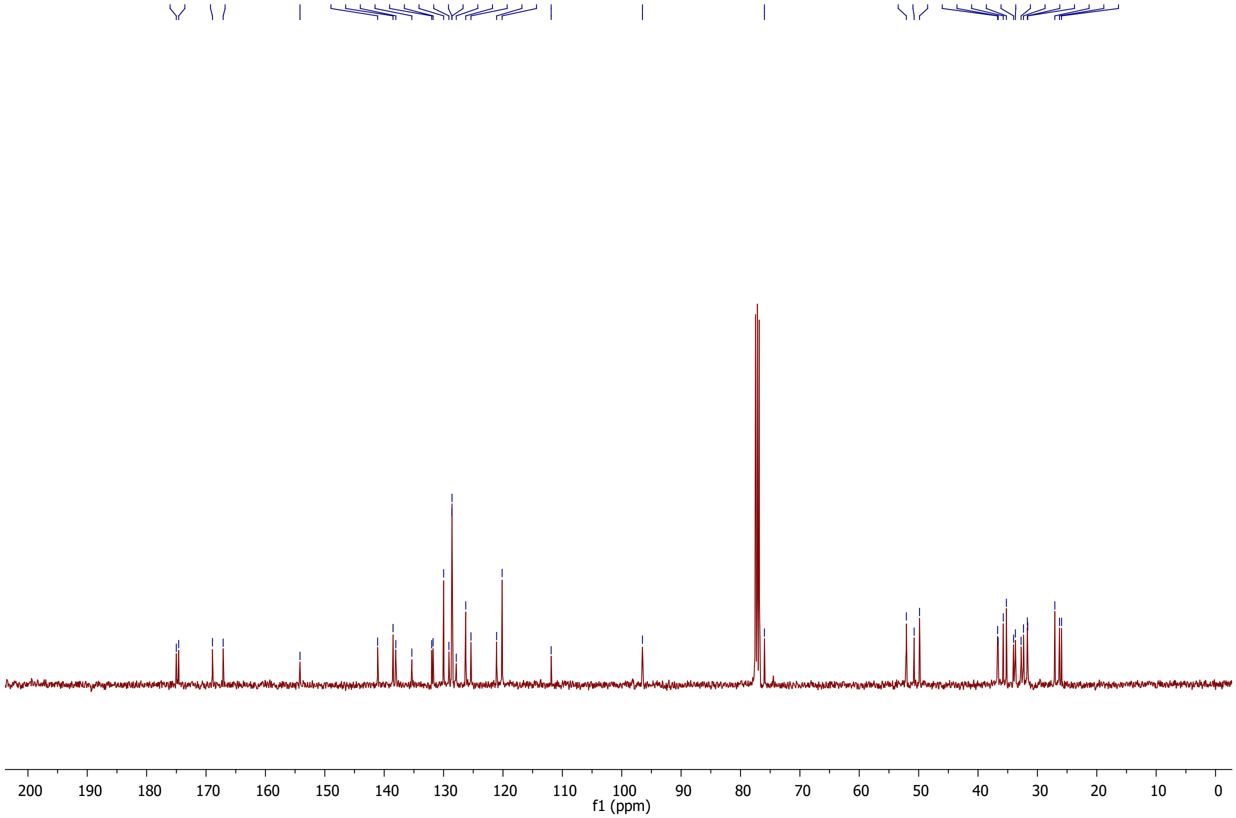

<u>MS of Probe **3** (ESI positive)</u>

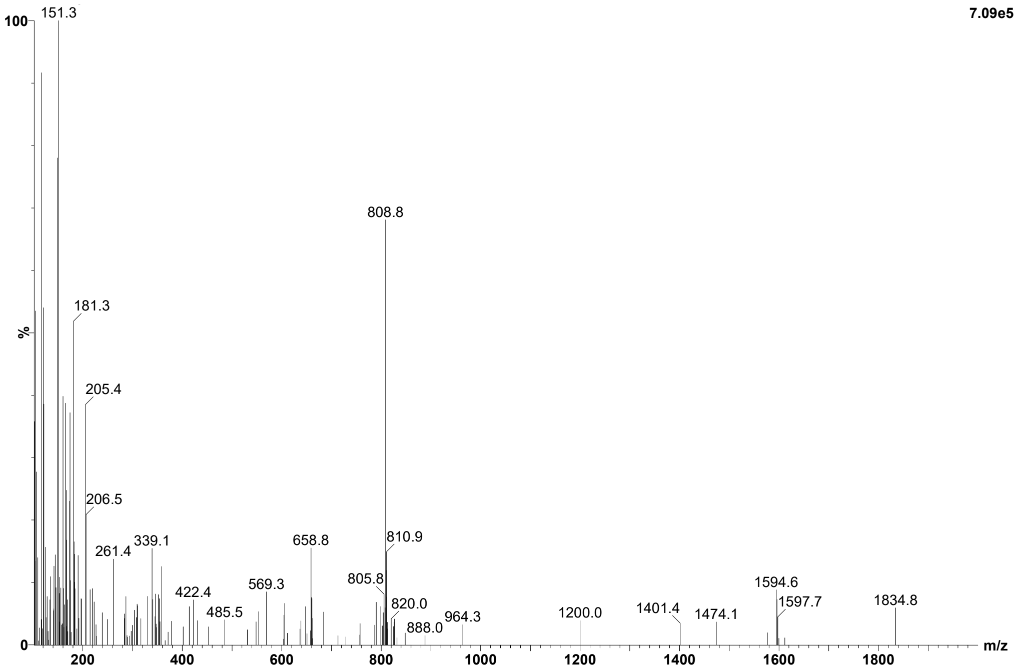

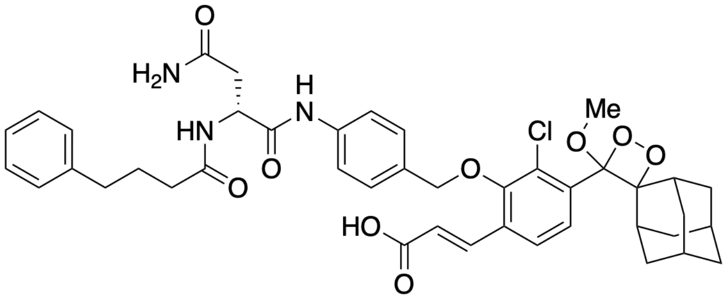

**Probe 4**

Chemical Formula: $C_{42}H_{46}ClN_3O_9$
Exact Mass: 771.29

[1]H NMR of Probe **4**

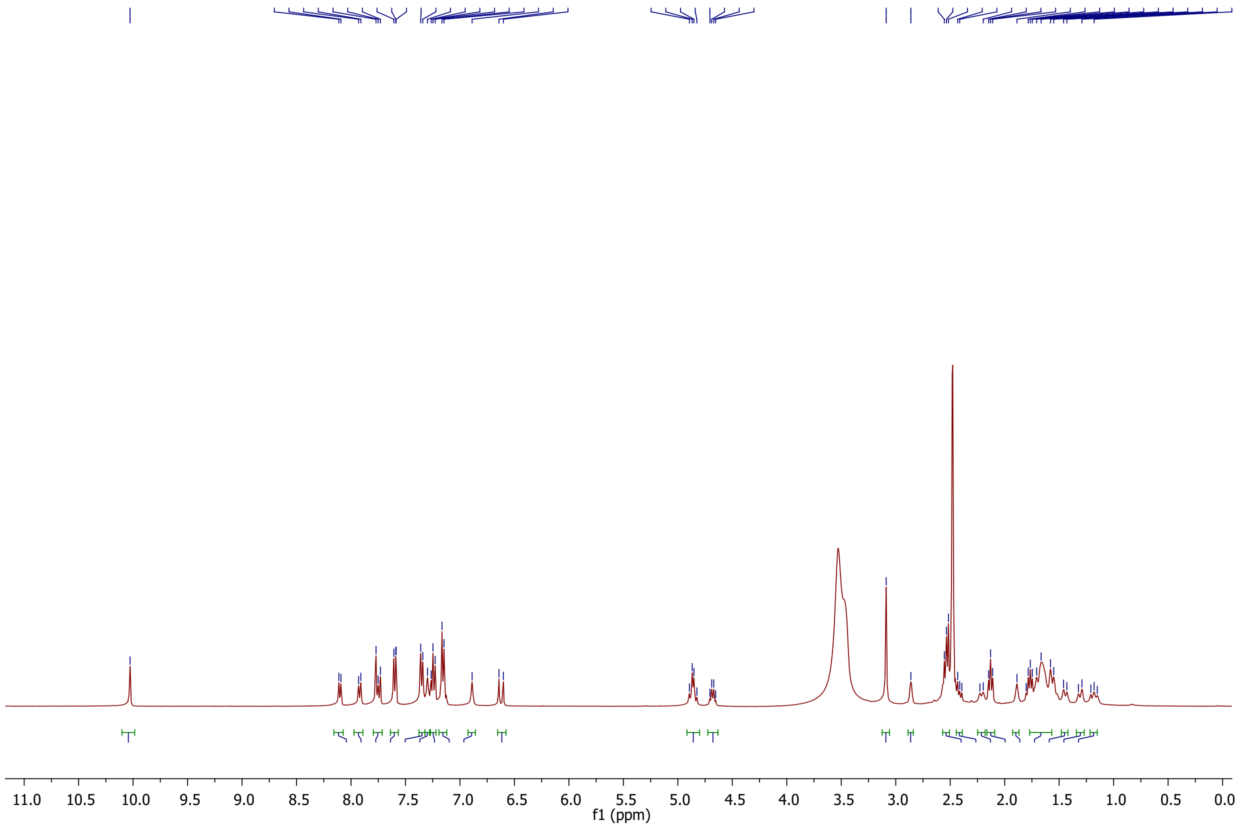

#### $^{13}$C NMR of Probe **4**

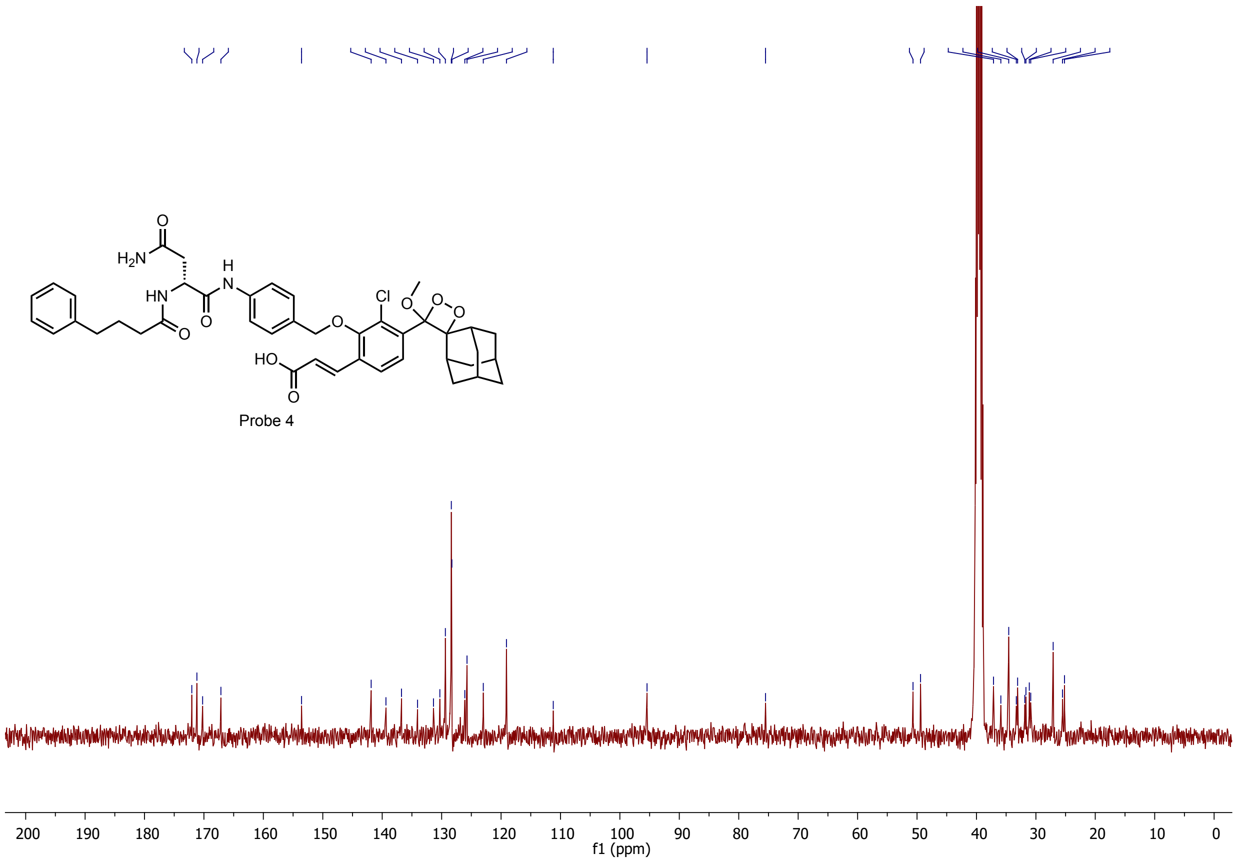

#### MS of Probe **4** (ESI negative)

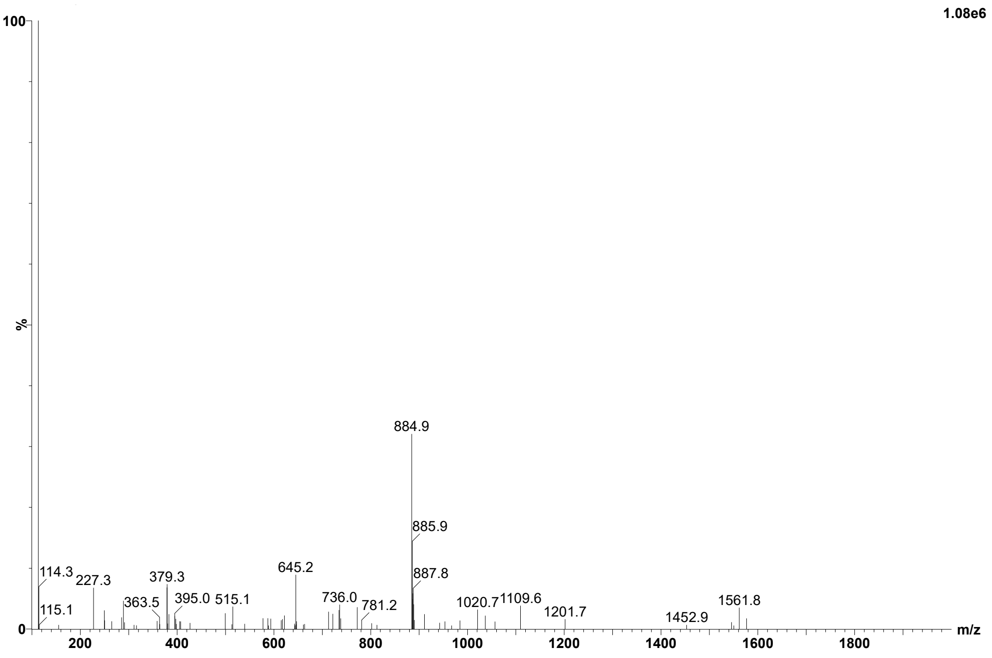

(8) Nougayrède, J.P.; Homburg, S.; Taieb, F.; Boury, M.; Brzuszkiewicz, E.; Gottschalk, G.; Buchrieser, C.; Hacker, J.; Dobrindt, U.; Oswald, E. Escherichia coli

induces DNA double-strand breaks in eukaryotic cells. *Science* **2006**, *313* (5788), 848–51.

(15) Wallenstein, A.; Rehm, N.; Brinkmann, M.; Selle, M.; Bossuet-Greif, N.; Sauer, D.; Bunk, B.; Spröer, C.; Wami, H.T.; Homburg, S.; von Bünau, R.; König, S.; Nougayrède, J.P.; Overmann, J.; Oswald, E.; Müller, R.; Dobrindt, U. ClbR Is the Key

Transcriptional Activator of Colibactin Gene Expression in *Escherichia coli*. *mSphere.* **2020**, *5* (4), e00591–20.

(16) Moretti, C.; Hosni, T.; Vandemeulebroecke, K.; Brady, C.; De Vos, P.; Buonaurio, R.; Cleenwerck, I. *Erwinia oleae sp. nov.*, isolated from olive knots caused by *Pseudomonas savastanoi pv. savastanoi*. *Int J Syst Evol Microbiol*. **2011**, *61* (Pt 11), 2745–2752.

(17) Shieh, W.Y.; Lin, Y.T.; Jean, W.D. *Pseudovibrio denitrificans gen. nov.*, *sp. nov.*, a marine, facultatively anaerobic, fermentative bacterium capable of denitrification. *Int J Syst Evol Microbiol*. **2004***, 54* (Pt6), 2307–2312.

(18) Madeira, F.; Madhusoodanan, N.; Lee, J.; Eusebi, A.; Niewielska, A.; Tivey, A.R.N.; Lopez, R.; Butcher, S. The EMBL-EBI Job Dispatcher sequence analysis tools framework in 2024. *Nucleic Acids Res*. **2024**, *52* (W1), W521–W525.
